# NFIC-mediated CaSR endocytosis defines a hyperactive TOMM20^high^CHGA^high^ oxyphil cell state as a pathological driver of autonomous secondary hyperparathyroidism

**DOI:** 10.64898/2026.05.06.723114

**Authors:** Qi Yang, Junsong Liu, Yuanyuan Wang, Rong Zhao, Hexiang Li, Yao Yao, Chongwen Xu, Bo Kou, Meng Lei, Qian Zhao, Xin Chen, Honghui Li, Ruimin Zhao, Rongrong Cui, Meichen Wang, Mengdan Li, Xiaobao Yao, Yanxia Bai, Fada Xia, Shaoqiang Zhang, Xi Liu, Xinying Li, Peng Hou

## Abstract

Secondary hyperparathyroidism (SHPT) is a debilitating complication of chronic kidney disease. Its clinical management is frequently compromised by calcimimetic resistance, a refractory state primarily driven by the progressive downregulation of the membrane calcium-sensing receptor (CaSR). Despite the centrality of CaSR as a therapeutic target, the mechanisms governing its aberrant expression and membrane localization remain incompletely elucidated. Here, we generated a single-cell transcriptomic atlas of human parathyroid tissues from SHPT and primary hyperparathyroidism (PHPT) patients, uncovering a unique stromal-immune niche that is specifically induced by uremic stress in SHPT. Our data also observed a striking dissociation between CaSR mRNA and its abundance as a membrane protein in SHPT tissues. Pseudotime trajectory analysis showed a progressive decline in CaSR pathway activity and a concomitant increase in endo-lysosomal activity along the trajectory, terminating in pathological oxyphil cells as the endpoint of chief cell differentiation in SHPT. Mechanistically, Nuclear Factor I C (NFIC) transcriptionally activated clathrin light chain B (CLTB) and Ras-related protein Rab7a (RAB7A) to trigger clathrin-mediated endocytosis and lysosomal degradation of CaSR, thus reducing its membrane abundance. This degradative program was further validated by multiplex immunofluorescence in TOMM20^high^CHGA^high^ pathological oxyphil cells in human SHPT tissues. To translate these mechanistic findings into a clinically actionable strategy, we repurposed the clinically approved lysosomal inhibitor hydroxychloroquine (HCQ) to block this CaSR degradation pathway. In patient-derived xenograft (PDX) mouse models, co-administration of HCQ and cinacalcet acted synergistically to restore membrane CaSR expression, normalize serum PTH and calcium levels and suppress parathyroid tumor growth more effectively than monotherapies. Collectively, our single-cell-guided study identifies an NFIC-driven endo-lysosomal program as a previously unrecognized mechanism underlying CaSR downregulation and calcimimetic resistance in SHPT, and establishes HCQ repurposing as a clinically tractable therapeutic strategy for patients with refractory SHPT.

## Introduction

Secondary hyperparathyroidism (SHPT) affects 30–50% of patients with end-stage chronic kidney disease (CKD) receiving dialysis (1, 2), characterized by diffuse-to-nodular parathyroid hyperplasia and excessive secretion of parathyroid hormone (PTH) driven by disordered mineral metabolism, including hypocalcemia, hyperphosphatemia, and vitamin D deficiency (3). Uncontrolled SHPT develops to high-turnover bone disease and vascular calcification, and is strongly associated with increased cardiovascular mortality in CKD patients (4, 5). Although calcimimetics such as cinacalcet, an allosteric agonist of the calcium-sensing receptor (CaSR), reduce PTH levels by 30%–50% in early-stage SHPT (6), they frequently fail in patients with refractory nodular disease (7). Their utility is further limited by gastrointestinal intolerance, hypocalcemia, and rapid PTH rebound following withdrawal, leaving high-risk parathyroidectomy as the only option for many vulnerable patients (4).

SHPT initially responds to hypocalcemia but often progresses to autonomous hyperparathyroidism despite fluctuations in extracellular calcium ([Ca^2+^]_e_) status. This progression reflects parathyroid cellular reprogramming under chronic uremic stress, which manifests clinically as early diffuse hyperplasia with asymmetric multi-gland involvement (8), a hallmark of emerging epithelial heterogeneity. It ultimately progresses to refractory nodular lesions with enhanced 99mTc-sestamibi (MIBI) avidity, indicative of predominant clonal proliferation and distinct metabolic phenotypes, and culminates in persistent autonomous hyperfunction (9, 10). Collectively, these features demonstrate that acquisition of autonomy in advanced SHPT stems from this reprogramming, which drives intra-glandular heterogeneity and selective clonal dominance. However, the precise mechanisms underlying parathyroid cellular autonomy and calcimimetic resistance are inadequately comprehended.

Histologically, this heterogeneity is further exhibited by significant pathological changes, particularly within calcimimetic-resistant nodules characterized by an elevated ratio of oxyphil cells to chief cells (11–13). Although oxyphil cells are generally considered mitochondrial-rich, senescent derivatives of chief cells in healthy parathyroid tissue, their prevalence in SHPT and primary hyperparathyroidism (PHPT) correlates positively with circulating PTH levels (14). This finding suggests that a certain subtype of oxyphil cells demonstrates hypersecretion and reduced sensitivity to calcium. However, current single-cell sequencing investigations have struggled to define a unique expression profile for oxyphil cells distinct from chief cells (15, 16); the discrepancy between their transcriptomic similarity and elusive pathogenesis highlights the complexity of the pathological mechanisms underlying SHPT.

CaSR serves as a primary regulator of parathyroid function. Membrane-localized CaSR senses [Ca^2+^]_e_ via G-protein-coupled pathways (Gi/o and Gq/11) to acutely suppress PTH secretion under physiological conditions (17–19). Additionally, CaSR suppresses parathyroid cell proliferation through β-arrestin/PKC/CaMK-mediated activation of ERK pathway (18). However, in SHPT, this “molecular brake” is functionally compromised. Augmenting CaSR sensitivity to [Ca^2+^]_e_ forms the pharmacological basis for calcimimetics. The repeated ineffectiveness of calcimimetics in treating advanced nodular hyperplasia suggests that current strategies are inadequate to address the intrinsic receptor deficits in heterogeneous glandular tissues. Thus, understanding the molecular mechanisms linking CaSR dysregulation to the emergence of pathological cellular phenotypes is imperative. Although previous studies have established CaSR downregulation as a hallmark of SHPT (7), with transcriptional repression driven by reduced vitamin D receptor (VDR) signaling, promoter hypermethylation, and dysregulated microRNAs partially explaining the diminished receptor abundance (20–22), the significance of CaSR expression heterogeneity on the development of epithelial autonomy and calcimimetic resistance remains largely unexplored.

Advanced SHPT exhibits convergent pathological charatceristics with PHPT, including autonomous cellular proliferation, excessive hormone secretion, resistance to physiological regulation by calcium and vitamin D, and nodular hyperplasia. However, in contrast to PHPT, which generally arises from genetic alterations, SHPT progression is uniquely driven by the chronic uremic stress. Therefore, PHPT serves as an ideal comparative model to dissect these complex pathogenic mechanisms. Here, we employed single-cell RNA sequencing (scRNA-seq) to systematically compare parathyroid tissues from patients with SHPT and PHPT. We aimed to isolate the specific molecular alterations induced by the uremic milieu, thereby delineating the mechanisms of SHPT progression and calcimimetic resistance to identify new therapeutic targets for this refractory disease.

## Results

### A high-resolution single-cell atlas defines the distinct immune-stromal ecosystem of SHPT

To elucidate the microenvironmental attributes of hyperparathyroidism (HPT) driven by distinct pathogenic mechanisms, we constructed a high-resolution single-cell transcriptomic atlas from 14 parathyroid specimens, comprising 1 case of parathyroid carcinoma (PC), 2 cases of parathyroid adenoma (PA), and 4 cases of SHPT (one classified as tertiary HPT post-transplantation; Figure 1A; Supplemental Table 1). After thorough quality control, normalization, and dimensionality reduction, the final dataset included 178,459 single cells (Figure 1B; Supplemental Figure 1). Parathyroid epithelial cells represented the majority of the cellular population in PC (80.25%), PA (89.36%), and SHPT (91.82%) tissues (Figure 1C-E; Supplemental Figure 2A, B), underscoring parenchymal proliferation as a shared pathological hallmark of HPT. However, distinct microenvironmental alterations were observed across the three groups. Specifically, PC tissues demonstrated a significant expansion of endothelial and immune cell populations compared to SHPT and PA (Figure 1E), an indication of the malignant features of sustained angiogenesis and tumor-promoting inflammation. Notably, a significant compositional imbalance was identified in this PC immune niche, characterized by a typical “myeloid-rich, lymphocyte-poor” phenotype (Figure 1F-H; Supplemental Figure 2C). The near-complete exclusion of T cells, natural killer (NK) cells, and dendritic cells (DCs) in PC signified a profound immunosuppressive barrier, highlighting the relative immune integrity of the SHPT/PA microenvironment.

**Figure 1.**
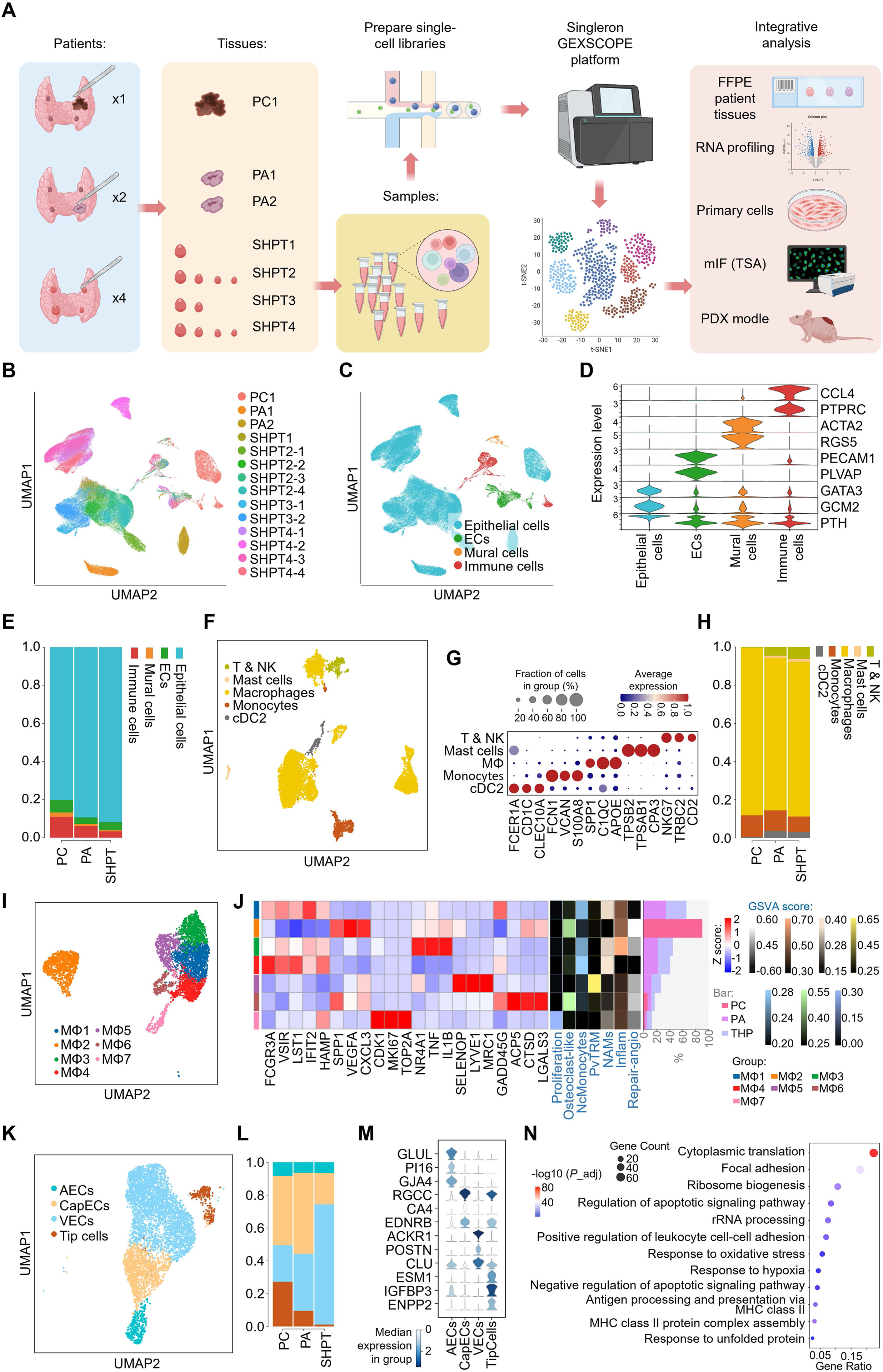
Single-cell transcriptomic landscape and cellular heterogeneity in parathyroid diseases. **(A)** Schematic workflow of the study. Parathyroid tissue samples were collected from patients with primary parathyroid carcinoma (PC, n = 1), primary parathyroid adenoma (PA, n = 2), and SHPT (n = 4) for single-cell RNA sequencing (scRNA-seq) using the Singleron GEXSCOPE platform, followed by integrative analysis and validation. **(B-C)** Uniform Manifold Approximation and Projection (UMAP) plots of all quality-control filtered cells, colored by sample origin **(B)** and major cell types **(C)**. **(D)** Violin plots showing the expression levels of canonical marker genes used to identify major cell lineages, including epithelial cells, ECs (Endothelial cells), mural cells, and immune cells. **(E)** Bar plot illustrating the proportional composition of major cell types across PC, PA, and SHPT groups. **(F)** UMAP visualization of the immune cell compartment re-clustered into five subtypes. **(G)** Dot plot showing the average expression and percentage of specific marker genes for each immune cell subtype. **(H)** The proportion of immune cell subtypes across the three disease groups. **(I)** UMAP plot showing the sub-clustering of macrophages (MΦ) into seven distinct clusters (MΦ1–MΦ7). **(J)** Heatmap displaying the expression of key functional genes and signature scores used for clarifying M1, M2a, M2c, proliferation, and tissue resident macrophages (TRM) across macrophage clusters. Histogram on the right side delineated the percentage of each cluster taking in the 3 disease groups. **(K)** UMAP visualization of ECs re-clustered into arterial (AECs), capillary (CapECs), venous (VECs), and Tip cells. **(L)** The cellular composition of EC subtypes across different groups. **(M)** Violin plots of marker genes identifying specific EC subpopulations. **(N)** Gene Ontology (GO) enrichment analysis of upregulated DEGs in SHPT-derived VECs from compared with PA- and PC-derived VECs.

In view of the myeloid dominance within the HPT immune ecosystem, we further categorized macrophages into 7 subclusters (Figure 1I; Supplemental Figure 3) and employed UCell scoring to analyze their functional diversity, including tissue residency and inflammatory stress. The PC-specific MΦ2 cluster possessed a distinct distribution on the UMAP projection, separating from SHPT- and PA-derived macrophages (Figure 1I). This cluster, characterized by high expression of *SPP1*, *TGFBI*, and *MSR1*, represents a distinct subset of SPP1^+^ tumor-associated macrophages (TAMs). These cells were likely educated by the malignant microenvironment to facilitate fibrosis and tumor progression (Figure 1J).

In stark contrast, two prominent populations of tissue-resident macrophages (TRMs) defined the ecological niche of SHPT and PA (Figure 1J). The most prevalent cluster, MΦ1, displayed a unique transcriptional profile (P2RY12^+^, CX3CR1^+^). This signature, which is similar to nerve-associated macrophages (NAMs) found in other peripheral organs (23), suggests its role as a resident monitor of local tissue homeostasis. MΦ5, on the other hand, represents a scavenger-like, perivascular-resident population (LYVE1^+^, FOLR2^+^, CD163^+^). In addition to these homeostatic populations, the local niche in SHPT/PA tissues included specific stress-associated macrophages. The MΦ3 population, which is conserved across both SHPT and PA, was identified by an inflammatory signature driven by immediate early genes (TNF^+^, IL1B^+^, NR4A1^+^). MΦ6 manifested as a ACP5^+^ osteoclast-like cluster. Rather than mediating distal bone resorption, this unique phenotype presumably reflects the power of HPT that drives systemic macrophages to osteoclasts development. We further identified FCGR3A^+^ non-classical monocytes (MΦ4), and a highly proliferative pool (MΦ7, MKI67^+^) responsible for local macrophage replenishment. Collectively, these findings contrast the resident-dominated, functionally diverse macrophage landscape in SHPT/PA with the monocyte-driven, immunosuppressive characteristic of PC.

The endothelial systems across the 3 groups also exhibited significant compositional differences. Specifically, we observed a progressive increase in the proportion of venous endothelial cells (VECs) in the order of PC < PA < SHPT, with VECs reaching a particularly high frequency (>75%) in SHPT. Conversely, arterial endothelial cells (AECs), capillary endothelial cells (CapECs), and tip cells (almost absent) were markedly compressed (Figure 1K-M; Supplemental Figure 4A, B), signifying a venous bias of the vascular network toward a low-pressure, high-permeability system. The expansion of VECs increased vascular density and shortens diffusion distances, thereby facilitating waste clearance, and the delivery of nutrients and oxygen to support the metabolic demands of cellular hyperplasia under chronic stress. Functionally, differentially expressed genes (DEGs) highly expressed in SHPT VECs were enriched in pathways related to translation and ribosome biogenesis, antigen presentation and T-cell activation, as well as stress response and anti-apoptosis (Figure 1N; Supplemental Figure 4C). These findings reflect an adaptive vascular remodeling process characterized by enhanced biosynthetic capacity and the acquisition of an immune sentinel function in SHPT VECs.

CellChat analysis uncovered a distinct intercellular communication landscape in SHPT compared to PA and PC (Supplemental Figure 5). Specifically, we observed robust, tissue-specific crosstalk between parathyroid epithelial cells and stroma cells in SHPT tissues. Parathyroid epithelial cells highly expressed the ligands CXCL12 and RARRES2, which targeted macrophages and monocytes expressing the cognate receptors CXCR4 and CMKLR1, respectively, indicating active immune recruitment. Conversely, macrophages served as the primary source of ligands TNFSF12, NRG2, and JAG1, which target to their receptors on parathyroid epithelial cells. These data delineate a unique paracrine feedback loop in SHPT wherein parathyroid epithelial cells actively recruit immune infiltrates, which in turn drive epithelial hyperplasia via paracrine signaling, a regulatory circuit that is notably absent in both PA and PC.

### Epithelial subtypes and CaSR expression heterogeneity in SHPT

Following batch correction, epithelial cells from PC, PA, and SHPT were stratified into six distinct subclusters (Figure 2A-C; Supplemental Figure 6). The PC-exclusive cluster Ep #6 (comprising 95% of PC epithelium) formed an isolated “oceanic island” on the UMAP projection, which was topologically distinct from epithelial cells of benign tissues (Figure 2A). Unlike the relative stable genome of Ep #1-5, this cluster exhibited extensive chromosomal copy number variations (CNVs) (Figure 2D, E; Supplemental Figure 7) and significant downregulation of the differentiation marker GCM2 (Figure 2F). Pathway enrichment analysis further revealed multi-category pathway enrichment and typical malignant features, including responses to oxygen levels (Figure 2G). Collectively, these genomic and transcriptomic signatures delineate Ep #6 as the malignant epithelial subpopulation.

**Figure 2.**
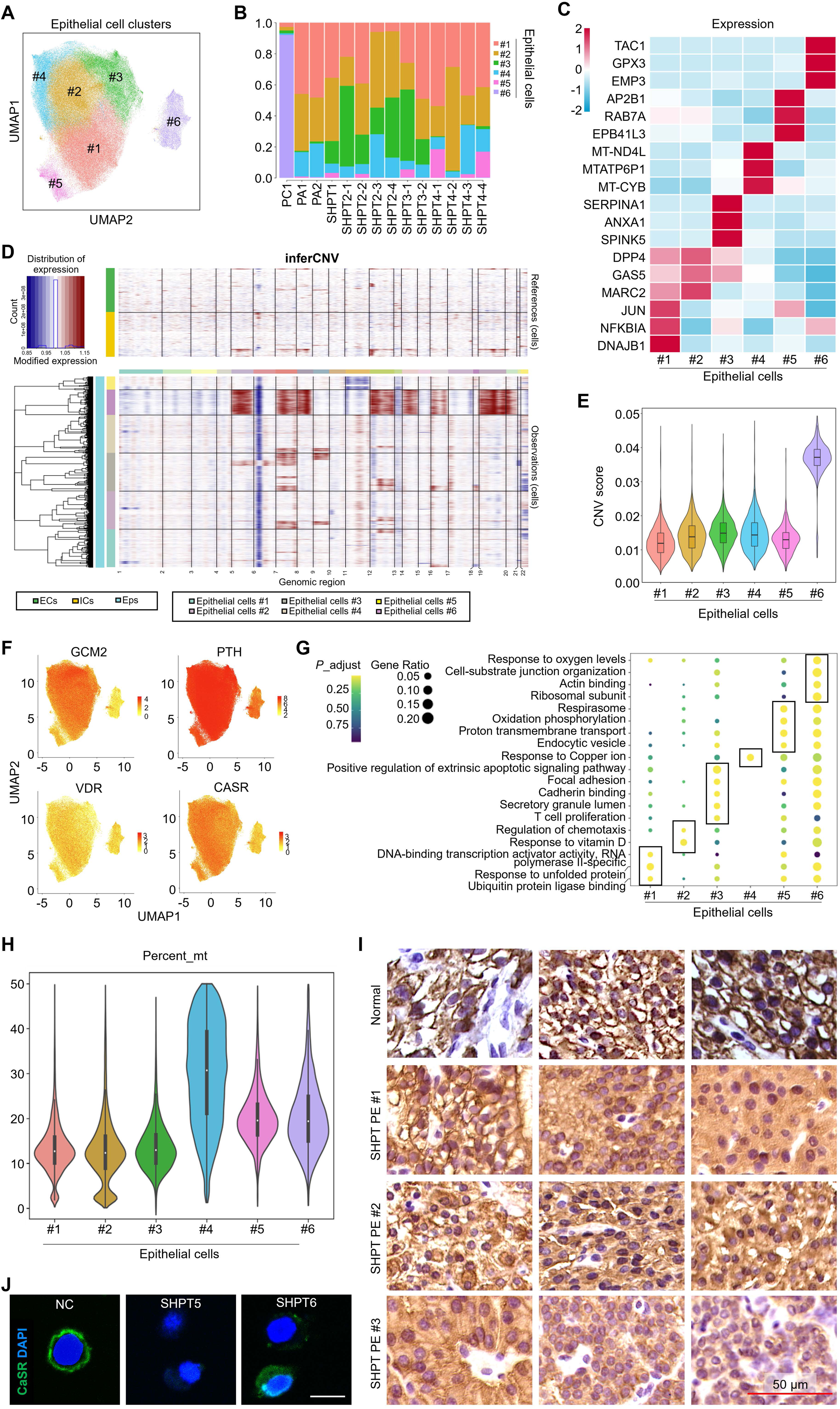
Transcriptomic heterogeneity of parathyroid epithelial cells and CaSR expression in SHPT cells. **(A)** UMAP visualization of re-clustered epithelial cells identifies six distinct sub-clusters (Ep #1–#6). **(B)** Stacked bar plot showing the proportional composition of the six epithelial clusters across individual samples. **(C)** Heatmap displaying the top differentially expressed marker genes specific to each epithelial sub-cluster. **(D-E)** Large-scale copy number variation (CNV) analysis inferred from scRNA-seq data. The heatmap (D) compares chromosomal alterations in epithelial clusters (Eps) against reference cells (ECs, endothelial cells; ICs, immune cells), and the violin plot (E) quantifies the CNV scores, highlighting significantly elevated genomic instability in Cluster #6. **(F)** Feature plots illustrating the expression distribution of canonical parathyroid functional markers (*GCM2, PTH, VDR,* and *CaSR*). **(G)** A dot plot of GO enrichment analysis that shows different biological pathways and functions that are linked to each epithelial cluster. **(H)** A violin plot that shows the percentage of mitochondrial gene expression (Percent_mt) across epithelial clusters. **(I)** Representative microscopic fields of immunohistochemistry (IHC) staining showing CaSR protein expression in normal parathyroid and SHPT tissue samples. Scale bar: 50 μm. (J) Immunofluorescence (IF) staining showing the cellular localization and expression of CaSR in primary parathyroid cells derived from normal and SHPT tissues. Green: CaSR; Blue: DAPI. Scale bar: 5 μm.

In contrast, clusters Ep #1–3 constituted the dominant epithelial bulk shared across all three groups (Figure 2A, B). These cells were enriched in pathways essential for stress response and cellular maintenance, including unfolded protein response (Ep #1), response to vitamin D (Ep #2), and cadherin binding (Ep #3) (Figure 2G). Furthermore, they displayed low mitochondrial transcript levels (Figure 2H). These transcriptional signatures are consistent with the functional baseline of parathyroid chief cells. Distinct from the chief cell lineage, Ep #4 and Ep #5 were characterized by significantly elevated mitochondrial content (Figure 2F). Ep #4 exhibited the most extreme mitochondrial enrichment and a unique “response to copper ion” transcriptional signature (Figure 2G). Ep #5, an SHPT-specific cluster, was individually distributed adjacent to the epithelial bulk population (Figure 2A, B). This cluster was more prevalent in the most severe SHPT case (SHPT4) (Figure 2B), and marked by enrichment of oxidative phosphorylation and respirasome pathways (Figure 2G). Given their high mitochondrial abundance and specific metabolic activity, we identified Ep #4 and Ep #5 as oxyphil cell subpopulations.

To characterize functional marker expression across these epithelial lineages, we found that PTH expression was universally high, while VDR, another key negative regulator of PTH secretion that is disrupted in early-stage SHPT, was broadly repressed (Figure 2F). Unexpectedly, CASR expression presented a considerable discrepancy from our cognition. Despite robust hypertranscription of *PTH*, *CASR* mRNA levels were maintained at moderate levels across all SHPT epithelial clusters. However, histological and cellular analyses revealed a profound defect at the protein levels. Unlike the membrane-localized CaSR enrichment observed in normal parathyroid tissues, SHPT tissues and primary parathyroid cells exhibited significant loss of CaSR membrane localization with cytoplasmic dissipating (Figure 2I, J). This discordance between maintained mRNA levels and defective protein localization delineates a critical post-translational dysregulation of CaSR that likely contributes to SHPT progression.

### Endocytic hyperactivity is a driver of SHPT pathogenesis

To decode the disease dynamics of HPT progression, we constructed a single-cell trajectory rooted in the quiescent chief cell cluster Ep #3, selected for its minimal transcriptomic diversity and low PTH secretory activity (Supplemental Figure 8). Pseudotime projection onto the UMAP plot visualized a progressive differentiation trajectory emanating from Ep #3 towards two distinct terminal states: the malignant Ep #6 and the SHPT-specific oxyphil cluster Ep #5 (Figure 3A-C). Branched Expression Analysis Modeling (BEAM) identified the molecular drivers underlying this fate bifurcation (Figure 3D). The pre-branch stage was defined by the upregulation of stress-response genes (*FOS*, *JUNB*, etc.; GO: response to unfolded proteins). Fate 2 (malignant trajectory) was driven by hypoxia and epithelial-mesenchymal transition (EMT) markers (*LDHA*, *VIM*, etc.; GO: response to hypoxia), whereas fate 1 (oxyphil trajectory) was characterized by the specific upregulation of endocytic regulators (*CLTB*, *RAB7A*, etc.; GO: clathrin-coated vesicle formation).

**Figure 3.**
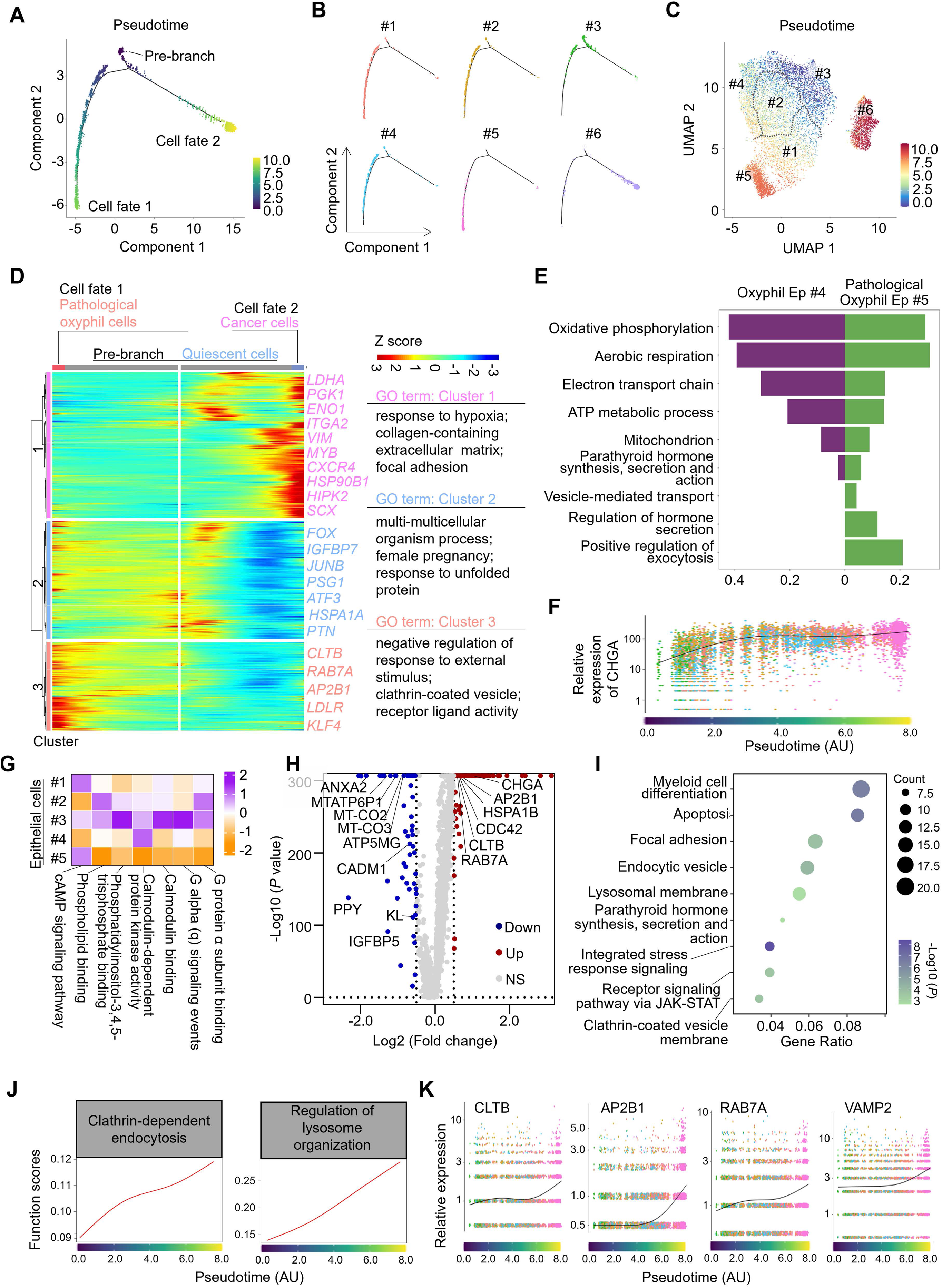
Distinct evolutionary fates of epithelial cells and functional divergence of pathological oxyphil cells. **(A)** Monocle 2 trajectory analysis inferring the developmental trajectory of epithelial cells, illustrating a progression from a “Pre-branch” state into two distinct cell fates (Cell fate 1 and Cell fate 2). The color scale represents pseudotime. **(B)** The distribution of the six epithelial sub-clusters along the inferred trajectory. **(C)** UMAP visualization overlaid with pseudotime values defined by Monocle 3 analysis. **(D)** Branched heatmap illustrating gene expression dynamics along the trajectory. Annotated GO terms highlighted functional characteristics of cells in different status. **(E)** Bar plot comparing pathway enrichment between oxyphil Ep #4 and pathological oxyphil Ep #5, highlighting their differences in hormone secretion activity. **(F)** Scatter plot showing the expression of the secretory marker *CHGA* along pseudotime. **(G)** Heatmap of GSVA scores for key signaling pathways downstream of CaSR across non-malignant epithelial clusters (Ep #1-#5). **(H)** Volcano plot displaying DEGs between Ep #5 and Ep #4. Red dots represent genes upregulated in Ep #5, while blue dots represent genes upregulated in Ep #4. (I) GO enrichment analysis of the upregulated DEGs in Ep #5. **(J)** Line plots showed the indicated pathway UCell score for cells ordered along pseudotime. **(K)** Pseudotime expression trends of key genes involved in endo-lysosomal pathway (*CLTB, AP2B1, RAB7A, VAMP2*).

This trajectory indicates a critical functional distinction between the two oxyphil populations (Supplemental Figure 9). In contrast to the ubiquitously distributed Ep #4, Ep #5 represented a terminal, SHPT-exclusive cellular state. Gene Set Enrichment Analysis (GSEA) provided a granular view of their functional divergence (Figure 3E). Both clusters exhibited robust enrichment in mitochondrial pathways (e.g., oxidative phosphorylation, aerobic respiration), reflecting their oxyphil nature, but they diverged sharply in secretory potential. Ep #5 uniquely acquired “peptide hormone secretion” capabilities, as evidenced high enrichment in pathways such as “positive regulation of exocytosis” and “regulation of hormone secretion”. This secretory phenotype was further supported by the highest expression of *CHGA*, a key regulator of secretory granule formation) (Figure 3F). Concurrently, Ep #5 showed a marked suppression of CaSR downstream signaling pathways such as calmodulin-dependent protein kinase activity (Figure 3G), suggesting minimal contribution of CaSR-coupled G protein signaling to the inhibition of PTH release. Additionally, genes upregulated in Ep #5 relative to Ep #4 were significantly enriched in “endocytic vesicle” and “lysosomal membrane” pathways (Figure 3H, I). Notably, UCell scores for “clathrin-dependent endocytosis” and “regulation of lysosome organization”, as well as the associated DEGs (*CLTB*, *RAB7A*, *AP2B1*, *VAMP2*), increased progressively along the pseudotime trajectory (Figure 3J, K). These data strongly suggest that Ep #5 is the pathogenic oxyphil subpopulation, where hyperactive clathrin-mediated endocytosis may drive disease progression by internalizing surface CaSR.

### Hyperactive clathrin-endosomal axis drives CaSR internalization and desensitization in SHPT

This proposed internalization mechanism implies a loss of surface CaSR function. Consistent with this, our aforementioned data have shown a gradual increase in clathrin-mediated endocytosis and lysosomal activity over the trajectory, accompanied by suppressed CaSR downstream signaling, such as “CaM kinase activity”, across multiple epithelial subclusters (Figure 3G, H). To validate this hypothesis, we isolated primary SHPT cells from different donors, and demonstrated that pharmacological inhibition of lysosomes with hydroxychloroquine (HCQ) or endocytosis with chlorpromazine significantly restored the membrane localization of CaSR in these cells (Figure 4A).

**Figure 4.**
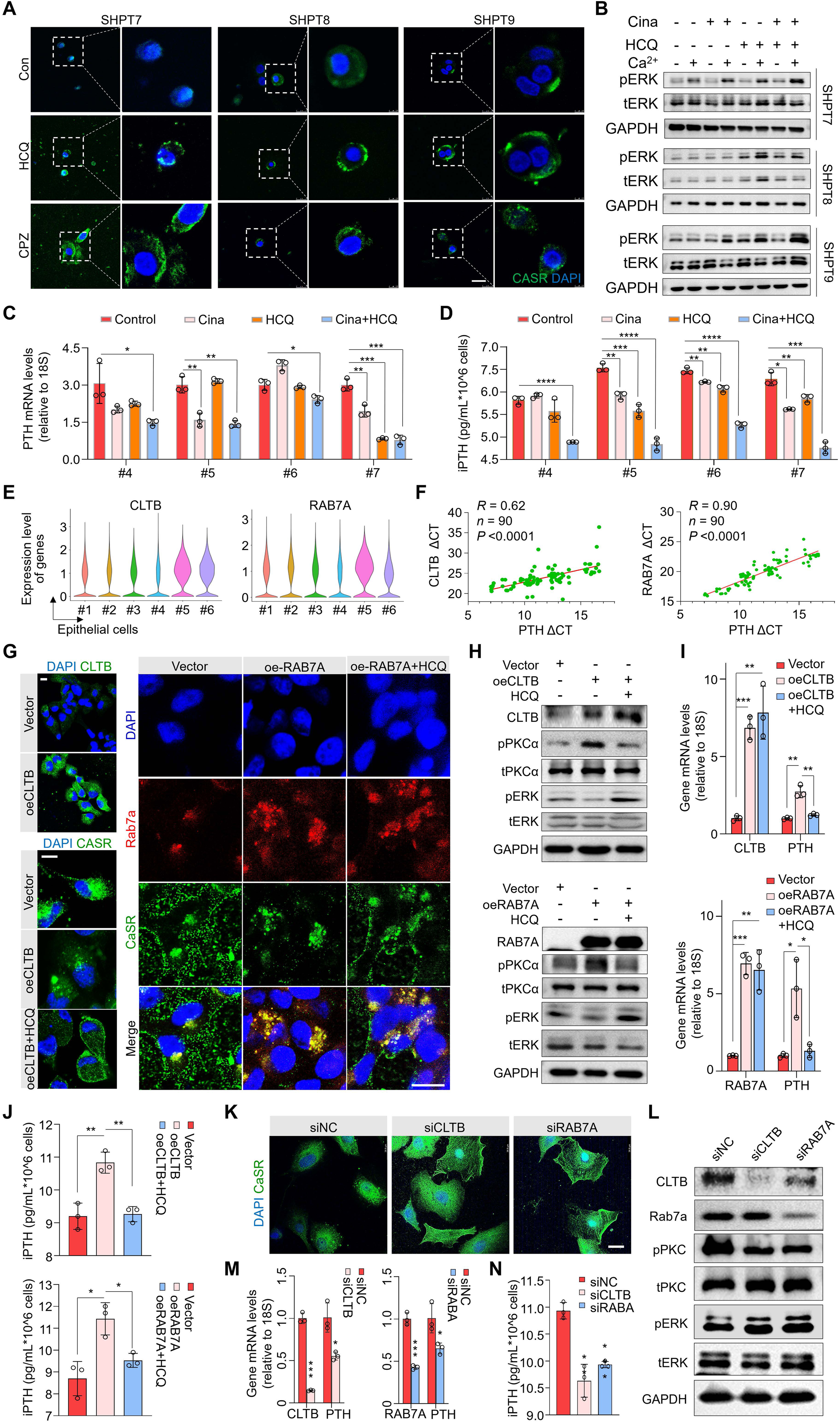
Modulation of CaSR localization and PTH secretion by *CLTB* and *RAB7A* in SHPT cells. **(A)** IF staining of CaSR (green) in primary SHPT cells treated with HCQ or CPZ, showing restored membrane expression. Scale bar: 5 μm. **(B)** Western blotting analysis of ERK phosphorylation (pERK) in SHPT primary cells that underwent the indicated treatments. HCQ, 20 μM HCQ treated for 24 h; Cina, 30 nM cinacalcet treated for 24 h; Ca^2+^, 2 mM calcium carbonate treated for 2 h. **(C-D)** Quantification of *PTH* mRNA levels (C) and secreted PTH concentration (D) under indicated drug treatments. **(E)** Violin plots illustrating the expression distribution of *CLTB* and *RAB7A* across epithelial sub-clusters based on our scRNA-seq data. **(F)** Quantitative detection of RNA extracted from SHPT paraffin-embedded tissues revealed a substantial positive link between *PTH* levels and vesicle genes (*CLTB*, *RAB7A*), as represented by ΔCT values. 18S rRNA (18S) served as the internal reference. **(G)** IF staining showing the expression and localization of CLTB, Rab7a, and CaSR proteins in primary SHPT cells following exogenous expression of *CLTB* (oe-CLTB) or *RAB7A* (oe-RAB7A) with or without HCQ treatment. Scale bar: 5 μm. **(H-J)** Cells in (G) were subjected to Western blotting analysis (H), showing the phosphorylation of PKCα (pPKCα) and ERK (pERK). GAPDH, total PKCα (tPKCα), and ERK (tERK) were served as internal references. Cells in (G) were subjected to mRNA detection (I) and iPTH examination (J). **(K-N)** SHPT primary cells were transfected with indicated siRNAs and subjected to IF staining against CaSR **(K)**, Western blotting analysis **(L)**, gene transcriptional quantification **(M)**, and iPTH detection **(N)**. Scale bar: 5 μm. 18S was used for normalization of mRNA levels. IPTH concentrations were normalized to cell numbers. Data are presented as mean ± SD. *, *P* < 0.05; **, *P* < 0.01; ***, *P* < 0.001.

Given that ERK is a canonical effector of CaSR signaling, its phosphorylation level serves as a reliable readout for CaSR sensitivity to [Ca²□]_e_ (18). While cinacalcet occasionally elevated the levels of phosphorylated ERK (pERK) in cultured SHPT cells, HCQ treatment, regardless of cinacalcet co-administration, significantly enhanced the responsiveness of CaSR to [Ca²□]e (Figure 4B). Furthermore, HCQ monotherapy reduced PTH transcription and secretion in primary cells from multiple SHPT patients, and its combination with cinacalcet maximized this inhibitory effect (Figure 4C, D). These findings, taken together, establish that targeting the clathrin-endosome axis can effectively reverse CaSR desensitization.

Based on the previous analyses of epithelial trajectories and DEGs analysis, we identified that CLTB and RAB7A were highly expressed in the pathological oxyphil cluster Ep #5 (Figure 4E). Moreover, RNA analysis of paraffin-embedded SHPT tissues showed a strong positive correlation between these genes and PTH expression (Figure 4F). Functionally, overexpression of either CLTB or RAB7A in primary cells abolished CaSR membrane localization, and this effect was effectively rescued by HCQ treatment (Figure 4G).

In parathyroid cells cultured in calcium-containing medium, exogenous expression of CLTB or RAB7A upregulated phosphorylated PKCα (pPKCα, Ser658) while downregulating pERK, an effect that was reversed by HCQ treatment (Figure 4H). These data indicate that CaSR depletion not only blunts canonical ERK signaling but also triggers aberrant, compensatory PKCα activation (24–26). Reflecting the above signaling perturbation, overexpression of CLTB or RAB7A promoted PTH transcription and secretion, and this pro-secretory effect was also abrogated by HCQ treatment (Figure 4H-J). Conversely, knockdown of CLTB or RAB7A restored CaSR membrane localization, suppressed pPKCα, enhanced pERK signaling, and reduced PTH transcription and secretion (Figure 4K-N). Taken together, these findings demonstrate that upregulation of CLTB and RAB7A accelerates CaSR endocytosis and lysosomal degradation, leading to receptor loss and signaling imbalance that drive excessive PTH secretion, thus highlighting this CLTB/RAB7A axis as a promising therapeutic target for restoring CaSR homeostasis.

### NFIC transactivates CLTB and RAB7A to drive CaSR endocytosis and degradation

To uncover the upstream driver of CLTB and RAB7A upregulation, we integrated regulatory network analysis (Figure 5A) with transcription factor (TF) binding site prediction (GTRD database) to screen for Ep #5-enriched TFs targeting the promoters of *CLTB* and *RAB7A*. Further analysis of the Cancer Cell Line Encyclopedia (CCLE) dataset (including 1,684 cell lines) revealed a significant positive correlation between *NFIC* expression and the expression of both *CLTB* and *RAB7A* (Figure 5B). Furthermore, GSEA revealed that genes co-expressed with *NFIC* were significantly enriched in endocytosis-related pathways (Figure 5C, D).

**Figure 5.**
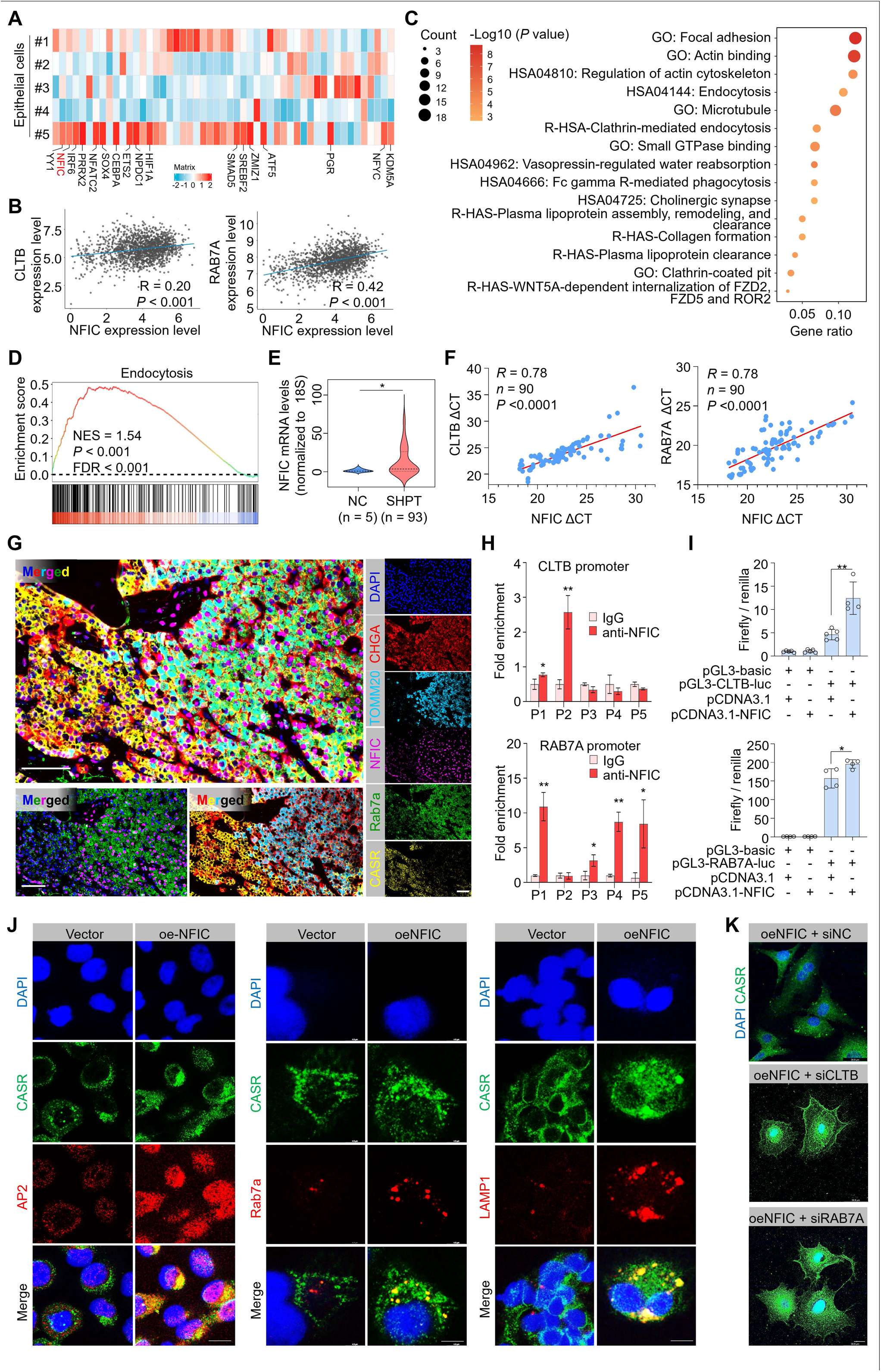
NFIC transcriptionally activates CLTB and RAB7A to regulate CaSR trafficking. **(A)** Heatmap displaying the expression profiles of transcription factors across nonmalignant epithelial cell clusters. **(B-D)** Correlation and functional enrichment analyses utilizing transcriptomic data from 1684 cell lines in the CCLE database. **(B)** Scatter plots demonstrated significant positive correlations between the expression levels of NFIC and CLTB or RAB7A. **(C)** GO/KEGG/Reactome pathway enrichment analysis and **(D)** GSEA of NFIC-correlated genes indicated significant enrichment in endocytosis-related pathways. **(E)** Violin plot showing upregulated NFIC mRNA levels in paraffin embedded SHPT tissues compared to normal controls (NC). **(F)** Correlation analysis revealing a significant positive association between NFIC and CLTB/RAB7A expression in in paraffin embedded SHPT tissues. **(G)** Multiplex IF staining of SHPT tissue sections showing the distribution of TOMM20 (mitochondria marker), CHGA (secretory granule marker), NFIC, Rab7a, and CaSR. Scale bars: 50 μm. **(H-K)** Mechanism investigations in SHPT primary cells. **(H)** ChIP-qPCR assays demonstrating direct binding of NFIC to specific promoter regions of *CLTB* and *RAB7A*. **(I)** Dual-luciferase reporter assays confirming that NFIC activates the transcription of *CLTB* and *RAB7A* genes. **(J)** IF images of SHPT cells overexpressing NFIC (oe-NFIC), showing increased expression and co-localization of CaSR with endocytic markers AP2, late endosomal marker Rab7a, and lysosomal marker LAMP1. **(K)** Rescue experiments showing that siRNA-mediated knockdown of CLTB or RAB7A (siCLTB/siRAB7A) abolished the CaSR endo-lysosomal accumulation induced by NFIC overexpression. Scale bars: 5 μm. Data are presented as mean ± SD. *, *P* < 0.05; **, *P* < 0.01.

Clinically, NFIC expression was markedly elevated in SHPT tissues compared to normal controls (Figure 5E), with its mRNA levels showing a strong positive correlation with those of *CLTB* and *RAB7A* (Figure 5F). Histologically, NFIC^high^ cells exhibited a TOMM20^high^CHGA^high^ signature (TOMM20, a mitochondria marker; CHGA a neuroendocrine marker), identifying them as the pathological oxyphil subpopulation defined in this study, distinct from both chief cell and conventional oxyphil cells (Figure 5G; Supplemental Figure 10A). Moreover, multiplex immunofluorescence (mIF) analysis showed that these pathological oxyphil cells displayed a Rab7a^high^CaSR^low^ profile (Figure 5G; Supplemental Figure 10B), while immunohistochemistry (IHC) staining on serial sections confirmed concurrent high CLTB expression in NFIC^high^ eosinophilic regions (Supplemental Figure 10C). These observations provide spatial evidence that NFIC orchestrates the in situ upregulation of this endocytic transport machinery.

Mechanistically, we confirmed the binding of NFIC to the promoters of *CLTB* and *RAB7A*, as well as its regulatory role in their transcriptional activity, using chromatin immunoprecipitation-quantitative PCR (ChIP-qPCR) and dual-luciferase reporter assays in primary SHPT cells (Figure 5H, I). As supported, NFIC overexpression substantially upregulated the expression of CLTB and Rab7a (Figure 5J; Supplemental Figure 11A), thereby causing the loss of surface CaSR, which co-localized with AP2 (a clathrin adaptor), Rab7a (a late endosome marker), and LAMP1 (a lysosome marker) (Figure 5J). Conversely, NFIC knockdown effectively restored plasma membrane localization of CaSR (Figure 5K). Crucially, NFIC overexpression suppressed CaSR downstream signaling, as evidenced by increased pPKCα and decreased pERK, and markedly enhanced PTH transcription and secretion. These pathological effects were effectively abrogated by the concurrent knockdown of *CLTB* or *RAB7A* (Supplemental Figure 11B).

Summarizing the aforementioned findings, we delineate a novel mechanism underlying CaSR downregulation and calcimimetic resistance in SHPT (Figure 6). Specifically, in SHPT, elevated NFIC transcriptionally upregulates CLTB and Rab7a, sequentially activating the clathrin-mediated endocytic trafficking pathway. This axis enhances the lysosomal degradation of membrane-localized CaSR, consequently abrogating its inhibitory control over PTH transcription and secretion. Consequently, the loss of surface CaSR fuels aberrant parathyroid cell proliferation and confers resistance to calcimimetics, ultimately driving SHPT progression. These data indicate that targeting the endo-lysosomal pathway offers a compelling therapeutic strategy, either as a direct intervention or as a means to restore cinacalcet sensitivity.

**Figure 6.**
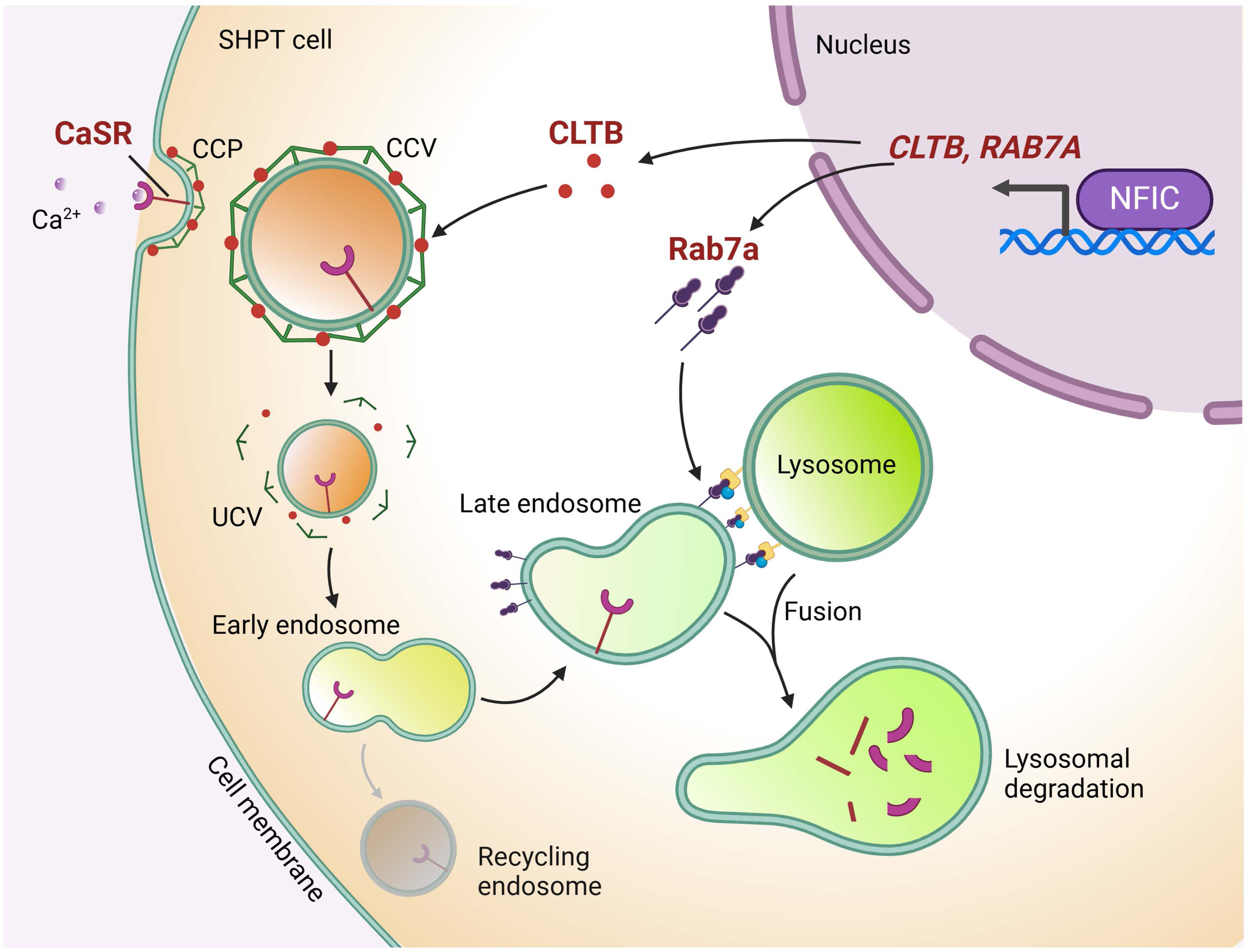
Schematic model of the NFIC-driven endocytic degradation of CaSR in refractory SHPT. Schematic illustration depicting how elevated NFIC orchestrates the downregulation of membrane CaSR in SHPT cells. NFIC acts as a master transcriptional regulator, simultaneously activating the expression of *CLTB* and *RAB7A*. This dual upregulation hyperactivates the clathrin-mediated endocytic pathway, where CLTB accelerates the internalization of surface CaSR into clathrin-coated vesicles (CCV), while Rab7a facilitates the trafficking of these vesicles through the endosomal system for fusion with lysosomes. This coordinated axis accelerates the trafficking of membrane CaSR toward lysosomal degradation rather than recycling. The resulting depletion of surface CaSR liberates the parathyroid cell from inhibitory control, fueling unrestrained PTH secretion, proliferation, and calcimimetic resistance.

### Therapeutic potential of HCQ in attenuating SHPT progression

Based on our *in vitro* findings that aberrant clathrin-mediated endocytosis drives CaSR desensitization and hyper-hormonal activity, we hypothesize that targeting this pathway may render a viable therapeutic strategy. To validate this *in vivo*, we established PDX models via subcutaneous implantation of freshly resected parathyroid tissues from 5 unrelated SHPT patients into immunocompromised nude mice. These recipient mice were then subjected to a 4□week treatment regimen to assess the therapeutic efficacy of HCQ alone or in combination with cinacalcet (Figure 7A). Longitudinal monitoring of serum human intact parathyroid hormone (iPTH) showed a sustained elevation in vehicle-treated mice post-transplantation, with a subset of PDX grafts displaying resistance to cinacalcet monotherapy (Figure 7B). In contrast, iPTH levels in the HCQ monotherapy and combination treatment groups started to decline after 2-3 weeks of intervention and reached a nadir by study endpoint (Figure 7B). Statistical analysis confirmed that iPTH levels were markedly lower in the combination treatment group relative to both vehicle□ and cinacalcet□treated groups. Concomitantly, both HCQ monotherapy and combination regimen significantly reduced serum calcium levels, while no significant intergroup differences in serum phosphorus were observed across all treatment groups (Figure 7C).

**Figure 7.**
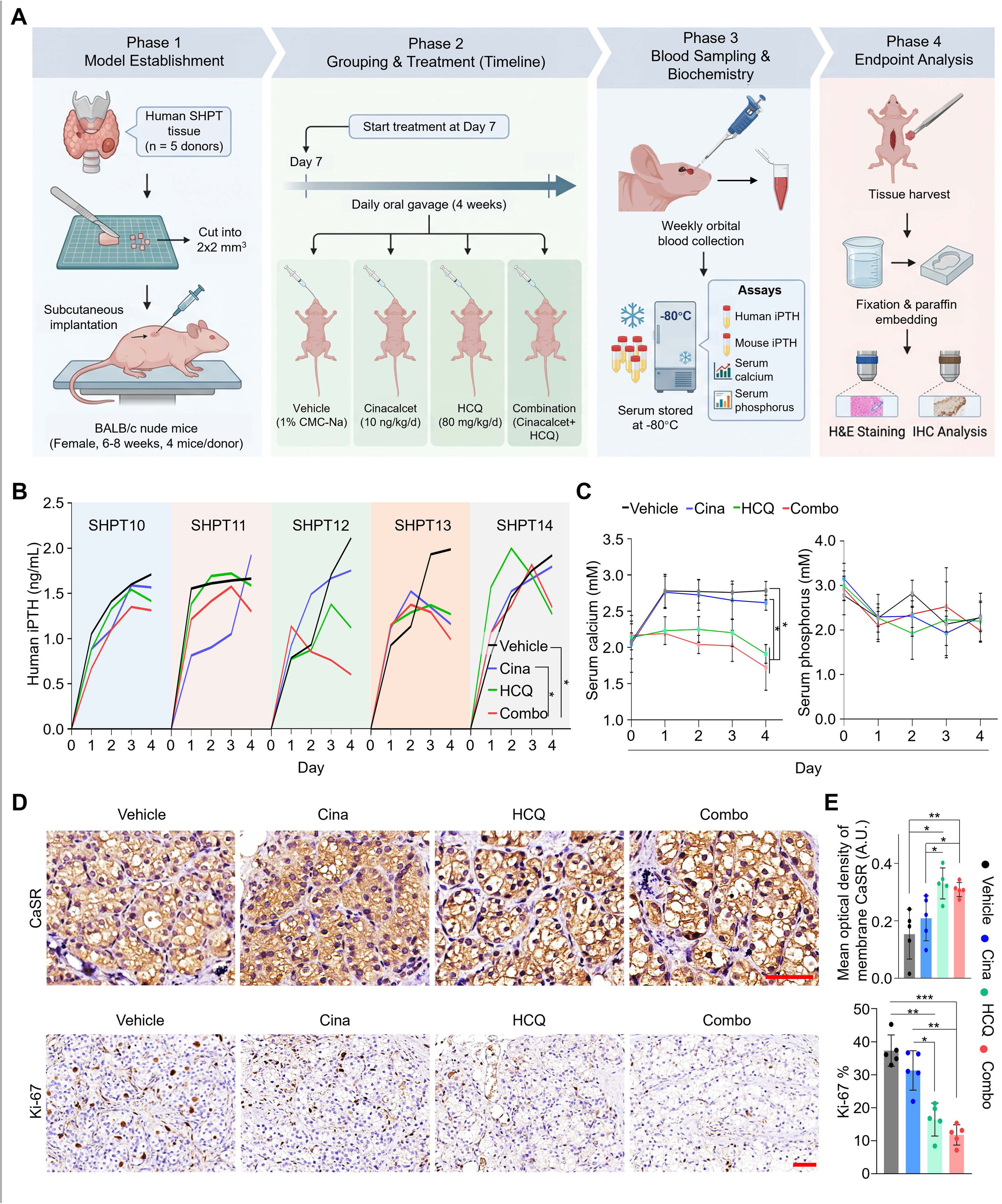
HCQ synergizes with cinacalcet to alleviate SHPT progression in a PDX mouse model. **(A)** Schematic diagram illustrating the experimental design of the PDX model. Briefly, human SHPT tissue fragments were subcutaneously implanted into nude mice. The established model mice were randomized into four groups: vehicle, cinacalcet, HCQ, or combination. Treatment was administered by daily oral gavage for 4 weeks, followed by serum biochemistry analysis and tissue harvesting. **(B)** Dynamic changes in serum human iPTH levels over the 4-week treatment period for five individual donors (SHPT10 - 14). **(C)** Time-course analysis of serum calcium (left) and serum phosphorus (right) levels throughout the treatment duration. **(D)** Representative IHC staining images of CaSR (top row) and Ki67 (bottom row) in graft tissues from the indicated groups. Scale bars: 50 μm. **(E)** Quantitative analysis of membrane CaSR mean optical density (top) and the percentage of Ki67-positive cells (bottom) across the groups. Cina, cinacalcet; Combo, combination. Data are presented as mean ± SD. *, *P* < 0.05; **, *P* < 0.01; ***, *P* < 0.001.

We next performed IHC staining of CaSR and proliferation marker Ki-67 in SHPT PDX xenografts across different treatment groups, and found that membrane-localized CaSR expression was strikingly restored in the HCQ monotherapy and combination treatment groups compared to vehicle- and cinacalcet-treated groups (Figure 7D, E, upper panel). By contrast, the nuclear positivity rate of Ki-67 was markedly reduced in mice receiving HCQ alone or combined with cinacalcet (Figure 7D, E, lower panel). Collectively, these findings indicate that HCQ effectively potentiates the therapeutic efficacy of cinacalcet by restoring membrane CaSR expression, thereby suppressing parathyroid cell proliferation and PTH secretion.

## Discussion

Cinacalcet resistance poses a major clinical challenge in the management of refractory SHPT (4, 7). Our study establishes that the substantial downregulation of membrane-localized CaSR constitutes the pivotal mechanism driving this disorder. Exploring the cellular roots of this deficiency, we observed substantial heterogeneity in CaSR expression within human SHPT tissues. Specifically, scRNA-seq identified a distinct population of pathological oxyphil cells characterized by near-complete loss of CaSR expression. Pseudotime trajectory analysis further provided unprecedented mechanistic insight: these pathological oxyphil cells represent the terminally differentiated endpoint of parathyroid cell differentiation. These observations imply that, during SHPT progression, parathyroid cells progressively transition towards this terminally differentiated state. More importantly, although aberrant activation of the endo-lysosomal pathway is initially identified in the oxyphil cell subpopulation by transcriptomic profiling, both bioinformatic interrogation and functional validation demonstrate that this pathogenic mechanism is broadly operative across all parathyroid cell types and SHPT tissues.

This widespread phenotypic transition likely reflects a state of cellular membrane remodeling triggered by chronic stress. The uremic milieu is characterized by excessive oxidative stress and accumulation of circulating toxins, including indoxyl sulfate and p-cresyl sulfate (27), which inflict continuous physical and chemical damage on the plasma membrane (28, 29). Parathyroid cells may adaptively enhance membrane turnover to preserve membrane integrity and cellular balance or to sequester incoming toxins. This survival-through-turnover approach has precedents in cell biology. Cells exposed to membrane damage from bacterial toxins swiftly activate endocytosis to eliminate damaged membrane segments (30). Likewise, renal tubular epithelial cells utilize a specialized program of membrane turnover and phagocytic engulfment to clear tubular luminal debris, thereby preventing harmful accumulation and sustaining organ function (31). We thus propose that, in the setting of SHPT, chronic exposure to uremic toxins forces parathyroid cells to engage a comparable defensive adaptation, in which CaSR internalization becomes collateral damage in the extensive membrane remodeling required to maintain homeostasis.

Furthermore, SHPT pathogenesis is not merely an isolated intracellular event but relies heavily on the specific tissue microenvironment. Through comparative analysis of SHPT and PHPT, we discerned unique stromal characteristics in SHPT, including a highly heterogeneous macrophage population and altered endothelial phenotypes, highlighting how systemic renal failure reshapes the local parathyroid niche to sustain a chronic inflammatory state. This exclusive epithelial-immune crosstalk in SHPT established a unique microenvironment that fuels inflammatory cell recruitment and inflammation□driven parathyroid cell proliferation. We thus postulate that sustained chronic stress, manifested as persistent inflammatory cytokine stimulation, oxidative stress, and uremic toxin accumulation, acts as the upstream trigger that induces aberrant overexpression of NFIC, thereby initiating the CLTB/Rab7a-mediated CaSR degradation pathway.

Upregulation of NFIC in SHPT aligns with the recognized role of the NFI family as stress-responsive remodeling factors. Precedents in chronic liver injury and cutaneous wound healing have established that NFI members are induced to orchestrate epithelial regeneration and structural repair (32–34). Moreover, in cancer biology, NFIB and NFIC have been demonstrated to directly regulate cell surface dynamics via modulating cytoskeletal organization and adhesion molecule expression (35, 36). Accordingly, we propose that NFIC upregulation in parathyroid cells represents a maladaptive response to chronic uremic stress. Although this process is presumed to preserve cellular homeostasis via enhanced membrane turnover, it ultimately compromises plasma membrane retention of CaSR.

Although acute CaSR activation is canonically established to induce phosphorylation of both PKCα and ERK, we noted an unanticipated inverse relationship between pPKCα and the CaSR-pERK axis upon modulation of the NFIC/CLTB/RAB7A pathway. This phenomenon may be attributed to the loss of CaSR triggering a local PTH elevation, which secondarily activates PKC via PTH1R (24). Additionally, PKCα signaling is usually controlled by strict negative feedback loops, such as RINCK-mediated degradation (25) and PHLPP-involved dephosphorylation (26). Dysfunction of such activation-induced downregulation may allow aberrant accumulation of pPKCα. This creates a vicious cycle as activated PKC phosphorylates CaSR at Thr888, which accelerates its internalization (37). Collectively, these observations indicate that dysregulated PKCα signaling constitutes a maladaptive response that exacerbates CaSR loss in SHPT, a process that warrants further mechanistic dissection.

Based on these mechanistic insights, we investigated the repurposing potential of HCQ as a therapeutic strategy for SHPT. Our findings support a conceptual shift in therapeutic design, a transition from high-dose calcimimetic-mediated activation of residual CaSRs to restoration of the functional receptor pool via inhibition of CaSR degradation. By suppressing lysosomal activity, HCQ effectively raises the amount of druggable CaSR on the cell surface. This multiplicative effect, in which HCQ restores CaSR abundance while cinacalcet enhances its activity, provides a molecular rationale for the therapeutic synergy observed in our models.

However, clinical translation of HCQ requires careful consideration and prudence. HCQ is predominantly eliminated through the renal route. SHPT patients typically present with uncorrected end-stage renal disease, where reduced clearance may result in drug accumulation, increasing the risk of severe adverse effects, including retinal damage (38). Moreover, HCQ is a pleiotropic agent with complex mechanisms; beyond lysosomal inhibition, it demonstrates significant immunomodulatory effects (39). The PDX mouse models employed in this study lack an integrated T-cell immune system, making it impossible to assess the potential impact of HCQ on SHPT pathogenesis via immune-mediated pathways. Consequently, while our findings with HCQ provides a proof-of-concept that refractory SHPT is reversible, future efforts should focus on the development of more selective endo□lysosomal inhibitors or the exploration of targeted local delivery strategies to realize therapeutic efficacy while minimizing safety concerns.

## Methods

### scRNA-seq data preprocessing and clustering

Computational analysis was performed using scanpy (v1.9.3) in Python(3.10). For each sample dataset, we filtered expression matrix by the following criteria: 1) cells with gene count < 200 or with top 2% gene count were excluded; 2) cells with top 2% UMI count were filtered out; 3) cells with mitochondrial content > 50% were removed; 4) genes expressed in less than 5 cells were excluded. Post-filtering, 178459 high-quality cells were retained. The raw count matrix was normalized by total counts per cell and logarithmically transformed into normalized data matrix. Top 2000 variable genes were selected by setting flavor = ‘seurat_v3’. Principle Component Analysis (PCA) was performed on the scaled variable gene matrix. The first 16 principle components (PCs) were used for clustering and dimensional reduction. Cells were separated into 28 clusters by using Louvain algorithm and setting resolution parameter at 1.2. Cell clusters were visualized by using Uniform Manifold Approximation and Projection (UMAP).

### Cell type annotation and differential expression analysis

To identify differentially expressed genes (DEGs), the scanpy.tl.rank_genes_groups function in scanpy was employed using the Wilcoxon rank sum test with default parameters. Genes were defined as DEGs if they met the following criteria: 1) detected in ≥ 10% of cells in either of the compared groups, 2) an average |log2FC | > 0.25 as DEGs, and 3) a Benjamini-Hochberg adjusted *P* value < 0.05. The canonical markers and their corresponding cell types were listed in Supplemental Table 2. Differentially expressed genes (DEGs) between specific groups or clusters were identified using the same criteria. Gene Ontology (GO), Kyoto Encyclopedia of Genes and Genomes (KEGG), and Reactome pathway enrichment analyses for DEGs were conducted using the clusterProfiler R package (v3.16.1) and ReactomePA R packages.

### Copy number variation (CNV) inference

To evaluate genomic instability and distinguish potential malignant epithelial cells, large-scale chromosomal CNVs were inferred from scRNA-seq data using the inferCNV R package (v1.8.0) (40). Endothelial and immune cells from the same dataset were used as normal diploid references. A cutoff of 0.1 was applied for the minimum average read counts per gene. The overall CNV burden for each cell was quantified as a CNV score, calculated as the sum of the squared absolute CNV values across all evaluated genes.

### Pseudotime trajectory analysis

To infer the developmental lineage and differentiation states of parathyroid epithelial cells, pseudotime trajectory analysis was performed using the Monocle 2 R package (v2.22.0) (41) and Monocle3 v1.0.0 (42). The top 2,000 highly variable genes were used to construct the trajectory using the DDRTree dimensionality reduction algorithm. Additionally, the CytoTRACE R package (v0.3.3) (43) was utilized to predict the differentiation potential (stemness) of single cells based on the number of expressed genes, providing an orthogonal assessment of cellular hierarchy.

### Gene set scoring and functional evaluation

To dissect the functional heterogeneity and estimate the relative activity of specific biological pathways across different cell clusters, Gene Set Variation Analysis (GSVA) was performed using the GSVA R package (v1.44.2) (44). For the assessment of macrophage polarization states and functional phenotypes, we utilized curated marker gene sets representing distinct modules (e.g., tissue-resident, pro-inflammatory signatures, and proliferation; detailed in Supplemental Table 3). Furthermore, to evaluate the functional divergence among epithelial subclusters (e.g., secretory and metabolic activities), we applied canonical pathway datasets, including hallmark pathways from the Molecular Signatures Database (MSigDB) and selected KEGG pathways. The resulting GSVA enrichment scores were utilized for comparative analysis and heatmap visualization.

### Cell-cell communication analysis

To infer intercellular communication networks within the parathyroid microenvironment, we utilized the CellChat R package (v1.6.1) (45). The analysis was based on the normalized expression matrix and the curated CellChatDB of human ligand-receptor interactions. We calculated the communication probability and interaction strength between major cell types and specific subclusters across different disease groups (PC, PA, and SHPT) using default parameters.

### Primary cell culture and treatments

Patient-derived parathyroid cell lines were established via primary culture of human parathyroid cell suspensions as previously described (46). A normal parathyroid cell line was derived from histologically normal parathyroid tissue adjacent to a parathyroid adenoma. These cells were grown in high-glucose DMEM medium supplemented with 10% fetal bovine serum (FBS), along with ampicillin and streptomycin. Cells were routinely passaged, cryopreserved, and used for drug treatment, ectopic gene expression, and gene silencing. All cell lines were periodically authenticated by STR profiling to exclude cross□contamination.

For drug treatments, cells were treated with 30 nM cinacalcet (catalog HY-70037, MCE), 20 μM lysosome inhibitor hydroxychloroquine (HCQ) (catalog T9287, TargetMol) or endocytosis inhibitor chlorpromazine hydrochloride (CPZ) (catalog HY-B0407A, MCE) for the indicated durations. DMSO served as the vehicle control. Where indicated, cells were substitutivity cultured in calcium-free high-glucose DMEM (catalog LA10096, Biolab, Beijing, China). In parallel experimental groups, 2 mM calcium carbonate was incorporated for a duration of 2 h.

For gene manipulation experiments, plasmids such as pCMV3-CLTB (NM_001834.3), pCMV3-RAB7A-Flag (NM_004637.5), and pCMV-3×HA-NFIC-Neo (NM_001245002.2) (1 μg per 1×10^5^ cells), or gene-specific siRNAs (25 pmol per 1×10^5^ cells) (Supplemental Table 4) were transfected into cells using Lipofectamine 3000 according to the manufacturer’s instructions.

### Cell immunofluorescence

Cells were fixed with 4% PFA, blocked with 5% BSA, and incubated with primary antibodies at 4°C overnight, followed by incubation with corresponding fluorescently labeled secondary antibodies. The nuclei were counterstained with DAPI. Images were acquired using a Zeiss LSM 880 confocal microscope.

### Tissue section staining

Paraffin-embedded parathyroid tissue sections were used for H&E staining, immunohistochemistry (IHC), and Tyramide Signal Amplification (TSA) multiplex immunofluorescence (mIF) staining (catalog abs0166, Absin Bioscience). H&E staining results were the base of morphological evaluation.

Standard IHC protocols were performed. Following antigen retrieval (citrate buffer, pH 6.0), sections were incubated overnight with primary antibodies against target proteins (details in Supplemental Table 5), followed by HRP-conjugated secondary antibodies and DAB visualization. Slides were digitally scanned (SRETT DX 12, Hungary). For quantitative analysis, Fiji (ImageJ) software was utilized to perform color deconvolution for DAB signal separation. The mean optical density of specific cellular compartments (nuclear or membrane) was calculated using the formula: mean optical density = log10(255 / mean gray value).

For TSA mIF staining, tissue sections underwent the standard procedures for deparaffinization, antigen retrieval, and blocking. Sections were subjected iterative cycles of primary antibody incubation (Supplemental Table 6), HRP-conjugated secondary antibody application, and specific TSA fluorophore deposition. Antibody complexes were stripped between each cycle via heat treatment (91°C in citrate buffer). Nuclei were counterstained with DAPI, and whole-slide multispectral images were acquired using a ZEISS Axioscan 7 system.

### Western blotting

Total protein was isolated utilizing RIPA lysate buffer supplemented with protease and phosphatase inhibitors. 30 μg of protein was separated by 6-12% SDS-PAGE, transferred to a PVDF membrane, blocked with 5% BSA, and incubated with the primary antibody overnight at 4°C. The membranes were incubated with HRP-conjugated secondary antibodies and visualized using ECL detection (WBKLS0500, Millipore).

### RNA extraction, reverse transcription and RT-qPCR

Total RNA from cultured cells was extracted using TRIzol reagent (Takara), while FFPE tissue samples were processed with the FFPE RNA Extraction Kit (R6954, Omega Bio-tek). Reverse transcription to generate cDNA was performed with the PrimeScript™ RT Reagent Kit (Takara) following the manufacturer’s instructions. Quantitative real-time PCR (qRT-PCR) was conducted using TB Green® Premix Ex Taq™ II (Takara). *RPS18* (18S rRNA) served as the housekeeping gene for normalization of target gene expression. Reactions were performed in triplicates, with primer sequences detailed in Supplemental Table 6.

### ELISA detection of intact PTH (iPTH)

We harvested the supernatant from the primary cell culture and performed cell counting. Otherwise, the blood serum was collected from the mouse in PDX study. The iPTH concentration was measured using the Human I-PTH ELISA Kit (catalog E-EL-H6076, Elabscience) in accordance with the manufacturer’s protocols. Absorbance was recorded at 450 nm employing a BioTek Cytation 5 (Agilent) microplate reader. The iPTH concentrations derived from the standard curve were normalized to cell number.

### CCLE database analysis

Cancer Cell Line Encyclopedia (CCLE) RNA-seq data (Expression Public 25Q2, 1684 cell lines) were available in the DepMap portal (https://sites.broadinstitute.org/ccle/datasets). Spearman correlation analysis identified NFIC-correlated genes (corr > 0.5, FDR < 0.05) across cell lines. GO/KEGG/Reactome enrichment was then performed. Specific correlations (NFIC *vs.* CLTB, NFIC *vs.* RAB7A) and GSEA were conducted using ggplot2 and clusterProfiler R package in R v4.4.0.

### Chromatin immunoprecipitation-qPCR (ChIP-qPCR)

ChIP-qPCR was performed using SimpleChIP® Enzymatic Chromatin IP Kit (CST) to evaluate NFIC enrichment at *CLTB* or *RAB7A* promoters. Cells (∼4×10□) were crosslinked (1% formaldehyde), chromatin sheared to 300-500 bp, immunoprecipitated (1-10 μg anti-NFIC *vs.* IgG), and DNA quantified by qPCR (primers shown in Supplemental Table 6) (n = 3 replicates).

### Promoter luciferase assays

Promoter luciferase assays confirmed NFIC transactivation of CLTB/RAB7A. Reporter plasmids pGL3-Basic-CLTB-Luc and pGL3-Basic-RAB7A-Luc were generated using the 3,000 bp upstream sequences from the TSSs (Transcription Start Sites). Primary cells were co-transfected with luciferase reporters (pGL3-Basic as the control), pRL-TK (normalization), pCMV-3×HA-NFIC-Neo or empty vector. After 48 h, lysates were assayed using Dual-Luciferase Reporter System (Promega) on BioTek Cytation 5 (Agilent). Data normalized as Firefly/Renilla ratios. n = 3 biological replicates.

### PDX models

Refractory SHPT tissues were obtained from surgically resected specimens (n = 5). Detailed clinicopathological characteristics of all patients are provided in Supplemental Table 1. One hundred mg (per implant) tissues were cut into fragments and implanted subcutaneously into the dorsal flank of 6–8-week-old female BALB/c nude mice using a trocar. Each patient□derived tissue was parallel inoculated into 4 mice. From day 7 post-implantation, mice were randomized to receive daily oral gavage of vehicle (1% CMC-Na), cinacalcet alone (10 mg/kg/day), HCQ alone (80 mg/kg/day), or the combination of cinacalcet and HCQ for 4 consecutive weeks. Blood samples were collected weekly from the orbital sinus before and after implantation. Serum samples were harvested and stored at −80°C for subsequent measurement of human iPTH (catalog E-EL-H6076, Elabscience), blood calcium (catalog ADS-W-D039, Aidisheng Biological Technology, Jiangsu, China), and blood phosphate (catalog ADS-W-LZ005, Aidisheng Biological Technology, Jiangsu, China). At the experimental endpoint, xenograft tissues were harvested, fixed, and paraffin-embedded for H&E and IHC staining following the standard protocols.

### Statistical analyses

Statistical analyses were performed using GraphPad Prism software (version 9.0.0, GraphPad Software). Data are presented as mean ± standard deviation (SD) unless otherwise stated. For comparisons between two groups (visualized as bar plots and violin plots), the unpaired or paired Student’s t-test was utilized. The correlation between gene expression levels was analyzed using Pearson correlation analysis. For time-course experiments involving repeated measurements (weekly changes in iPHT, blood calcium, and phosphorus), differences among groups were analyzed using a two-way analysis of variance (ANOVA), followed by multiple comparison tests (e.g., Tukey’s post hoc test) to determine statistical significance at specific time points. A *P* value < 0.05 was considered statistically significant.

### Study approval

The use of surgically resected tissues for this study was reviewed and approved by the Ethics Committee of The First Affiliated Hospital of Xi’an Jiaotong University (Approval No. LLSBPJ-2025-54). Broad written informed consent for the use of residual surgical tissues for medical research was obtained from all patients upon admission or prior to surgery, in accordance with the hospital’s standard policies for a teaching and research institution. The study was conducted in strict adherence to the principles of the Declaration of Helsinki. The animal study was approved and supervised by the Biomedial Ethics Committee of Health Science Center of Xi’an Jiaotong University (Approval No. XJTUAE2025-518).

## Data availability

The raw sequence data reported in this paper have been deposited in the Genome Sequence Archive for Human (GSA-Human) at the National Genomics Data Center, China National Center for Bioinformation / Beijing Institute of Genomics, Chinese Academy of Sciences (GSA-Human accession: HRA017650; BioProject accession: PRJCA061240). The dataset is publicly accessible at https://ngdc.cncb.ac.cn/gsa-human upon publication (47, 48). All other data supporting the findings of this study are available within the article and its Supplementary Materials.

## Author contributions

Conceptualization: P.H., Q.Y., and X.Li. Formal analysis: Q.Y., J.L., Y.W., He.L., Y.Y., R.C., S.Z., M.Li, and X.Liu. Funding acquisition: Q.Y. and M.W. Methodology: Q.Y., J.L., S.Z., X.Liu, X.Li, and P.H. Project administration: Q.Y., P.H., X.Li, X.Liu, and S.Z. Resources: J.L., C.X., B.K., M.Lei, Q.Z., X.C., Ho. L., R.Z., X.Y., Y.B., F.X., S.Z., X.Liu, X.Li, and P.H. Software: Y.Y. Supervision: Q.Y. and P.H. Validation: Q.Y. Visualization: Q.Y., He.L., Y.Y. Writing-original draft: Q.Y. and P.H. Writing-review and editing: all authors.

## Acknowledgments

We thank Drs. Jiao Fu, Hongjun Lv, Shu Liu and other colleagues from the Department of Endocrinology and Metabolism, the First Affiliated Hospital of Xi’an Jiaotong University, for their valuable advice and critical comments on this study. We extend our gratitude to Ang Su from Singleron Biotechnologies for his excellent support, which greatly facilitated this research.

