## Supplemental Figures for "NFIC-mediated CaSR endocytosis defines a hyperactive TOMM20^high^CHGA^high^ oxyphil cell state as a pathological driver of autonomous secondary hyperparathyroidism"

Qi Yang *et al*.

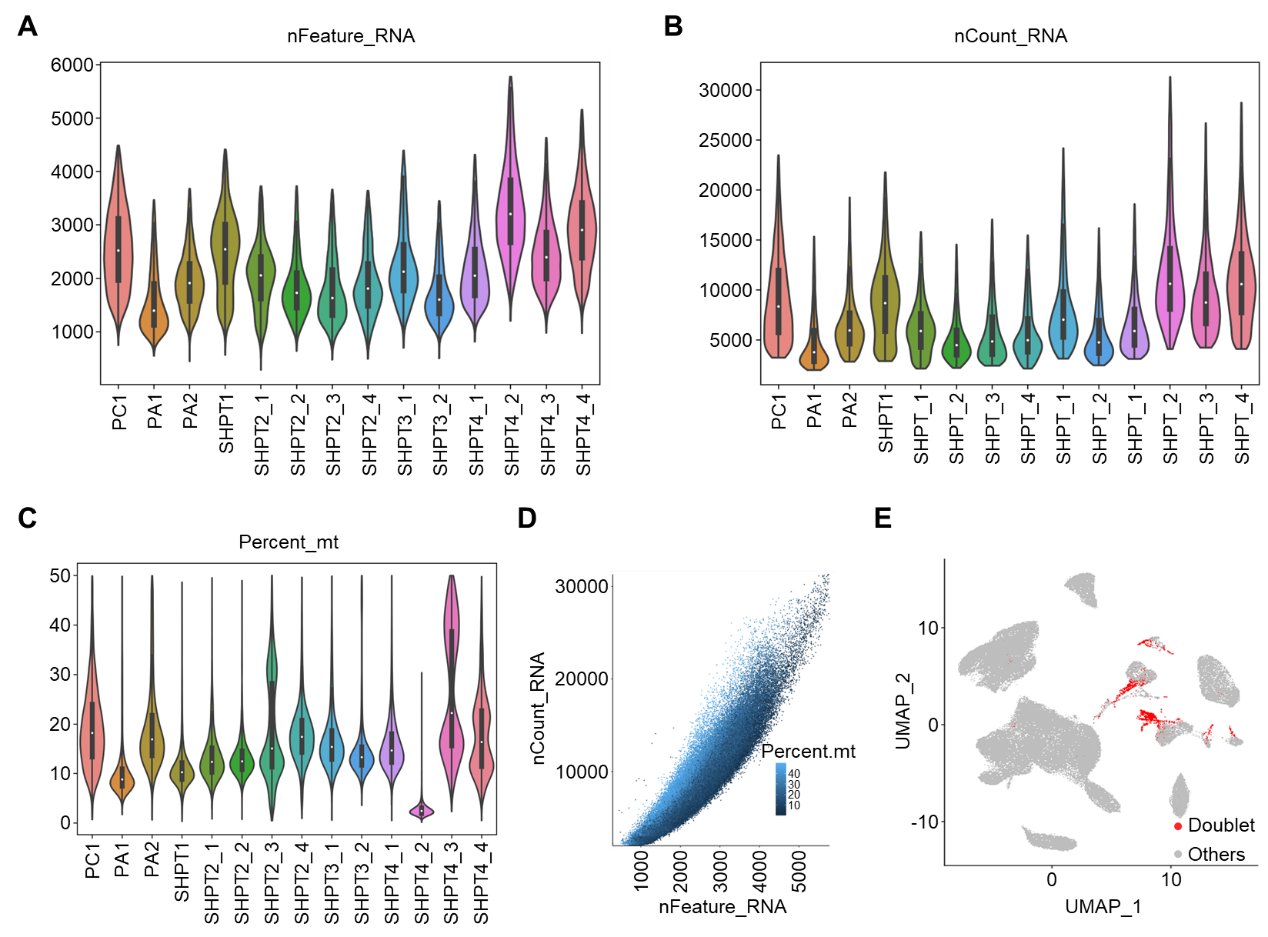
**Supplemental Figure 1. Quality control metrics and data pre-processing of single-cell RNA sequencing data.** **(A-C)** Violin plots showing the distribution of quality control metrics across different samples (PC, PA, and THP series). The panels display the number of detected genes per cell (**A**, nFeature_RNA), the total count of unique molecular identifiers (UMIs) per cell (**B,** nCount_RNA), and the percentage of mitochondrial gene expression (**C**, Percent_mt). **(D)** Scatter plot illustrating the correlation between sequencing depth (nCount_RNA) and the number of detected genes (nFeature_RNA). Each dot represents a single cell, colored by the percentage of mitochondrial reads. **(E)** UMAP visualization highlighting the identification of doublets. Predicted doublets are marked in red, while retained singlet cells are shown in gray.

**Supplemental Figure 2. Cellular composition and molecular characterization of the**
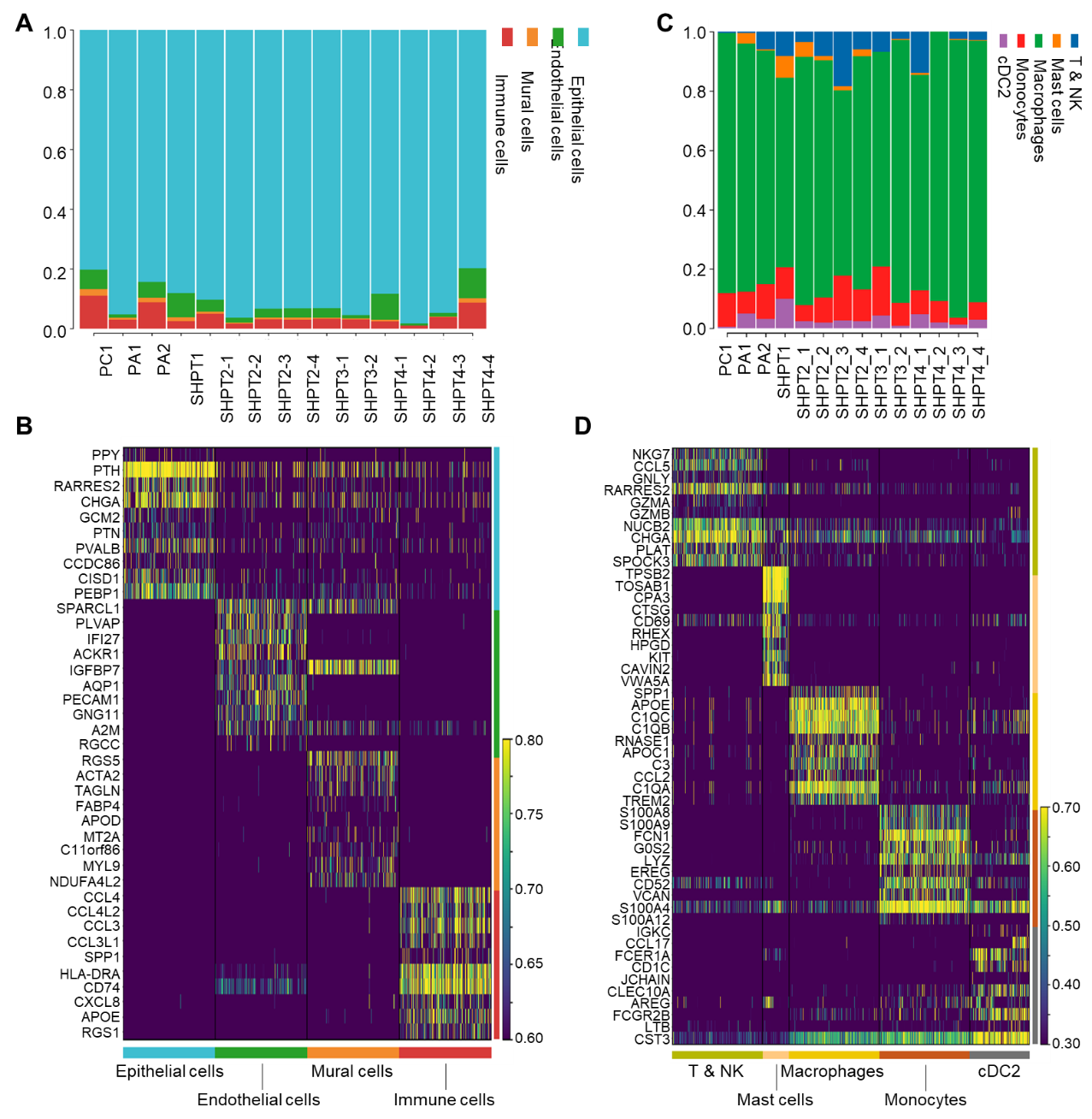
**parathyroid single-cell atlas. (A)** Stacked bar plot showing the relative proportions of the four major cell compartments (epithelial cells, endothelial cells, mural cells, and immune cells) across all 11 samples (PC, PA, and SHPT). **(B)** Heatmap displaying the scaled expression of the top 10 DEGs for each major cell type. **(C)** Stacked bar plot detailing the fractional composition of immune cell subtypes (T & NK cells, mast cells, macrophages, monocytes, and cDC2) within the immune compartment for each sample. **(D)** Heatmap showing the top 10 DEGs for distinct immune cell clusters.

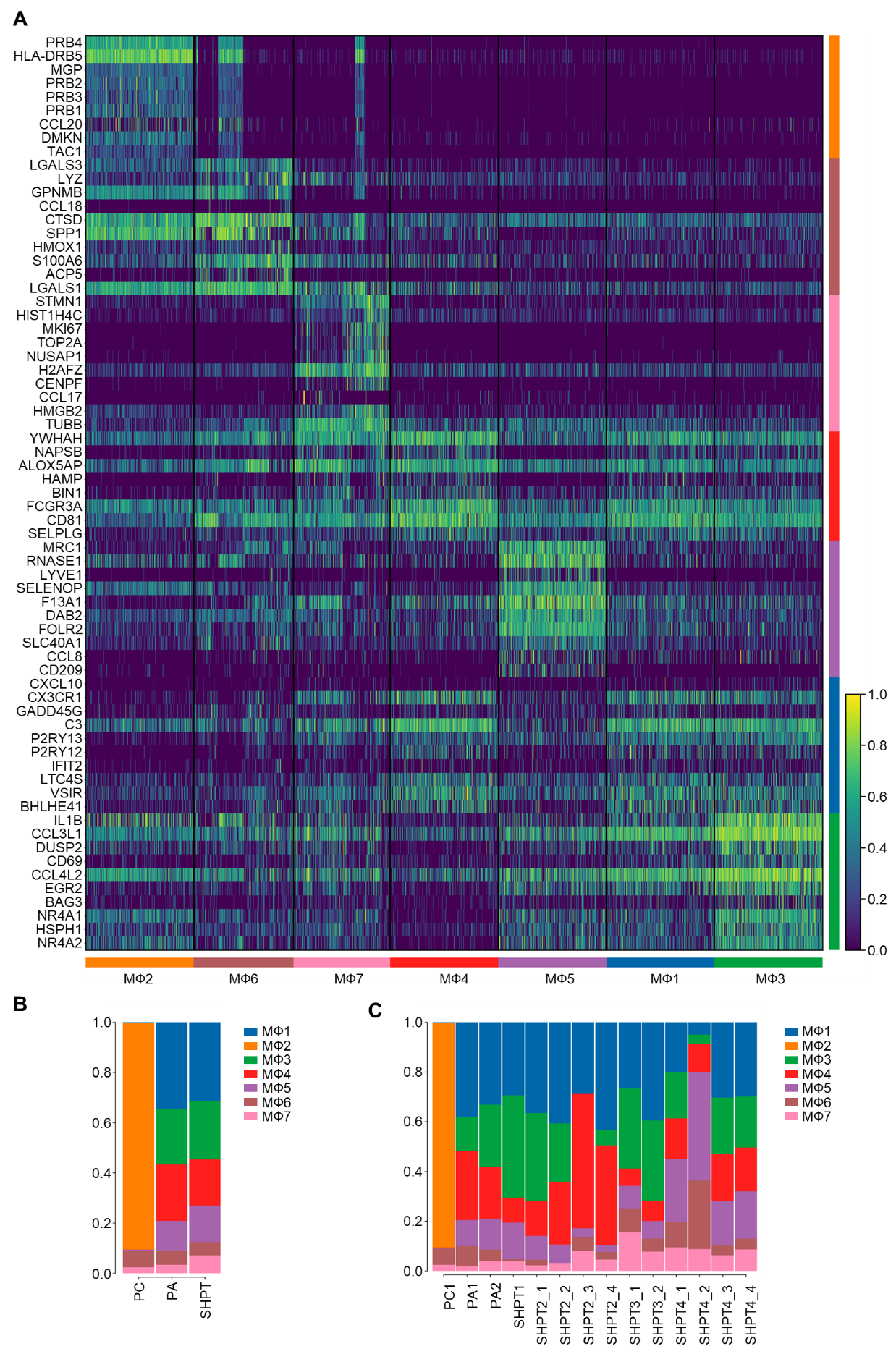
**Supplemental Figure 3. Transcriptomic signatures and proportional distribution of macrophage subpopulations.** **(A)** Heatmap displaying the scaled expression of top marker genes for the seven identified macrophage clusters (MΦ1–MΦ7). **(B)** Stacked bar plot showing the fractional composition of macrophage subpopulations across the three disease groups (PC, PA, and SHPT). **(C)** Stacked bar plot detailing the proportion of macrophage clusters within each individual sample.

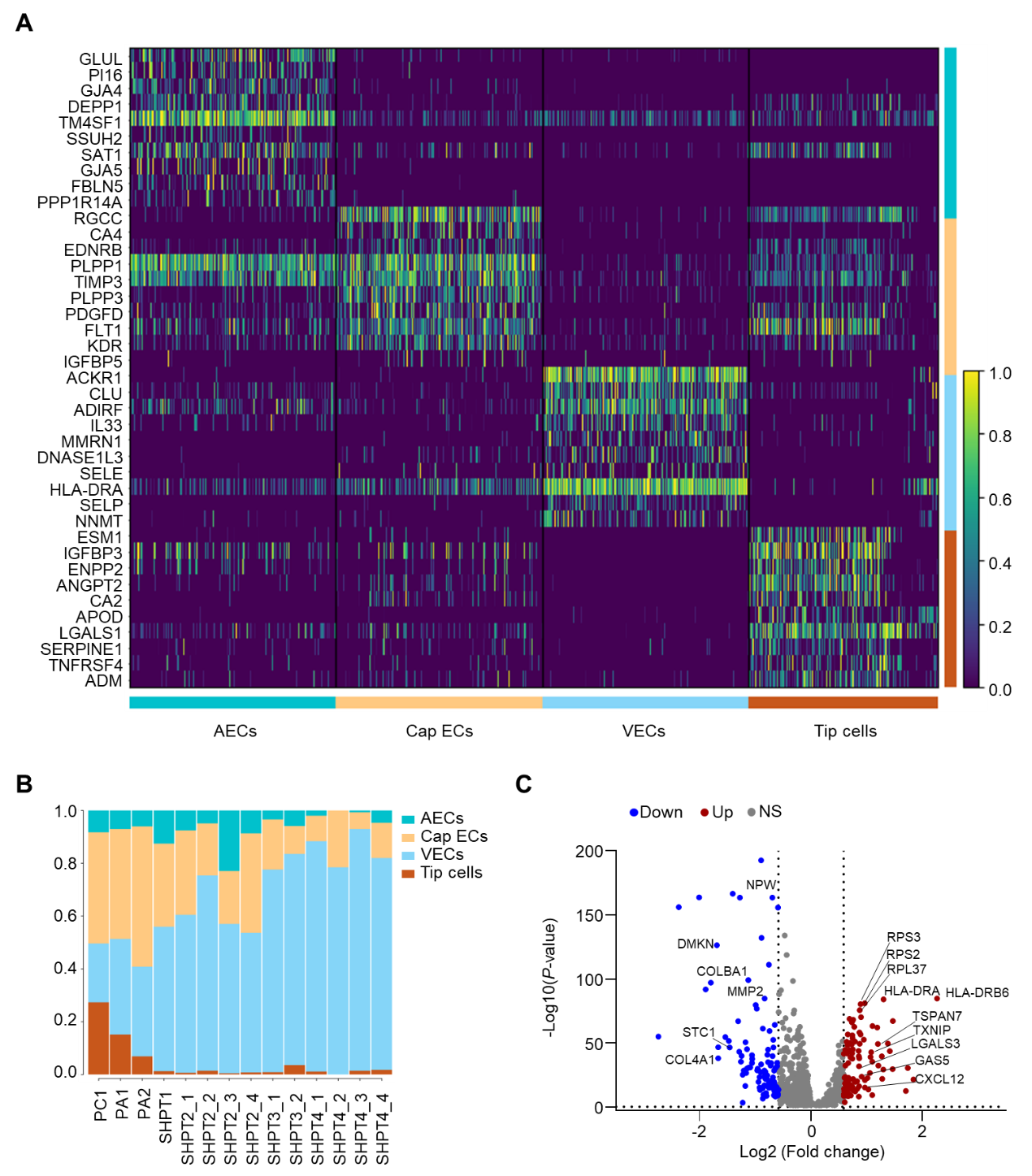

**Supplemental Figure 4. Characterization of parathyroid endothelial cell heterogeneity.** **(A)** Heatmap visualization of the DEGs defining four distinct endothelial subclusters. **(B)** Stacked bar plot illustrating the proportional distribution of the four endothelial subclusters across individual patient samples. **(C)** Volcano plot showing DEGs between VECs from SHPT vs. PC + PA. Red dots indicate significantly upregulated genes (log2FC > 0.585), and blue dots indicate downregulated genes (log2FC < -0.585).

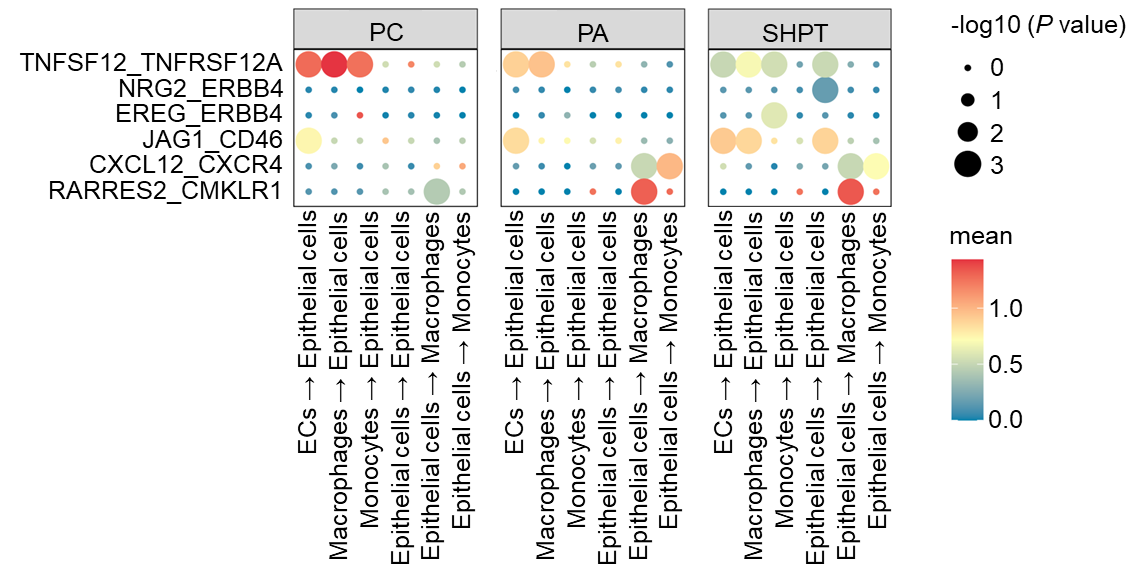

**Supplemental Figure 5. Comparative analysis of cell-cell communication networks in different parathyroid diseases.** Dot plots illustrating the significant ligand-receptor interactions between different cell populations in PC, PA, and SHPT tissues. The rows represent specific ligand-receptor pairs, while the columns indicate the interacting cell groups (sender→receiver).

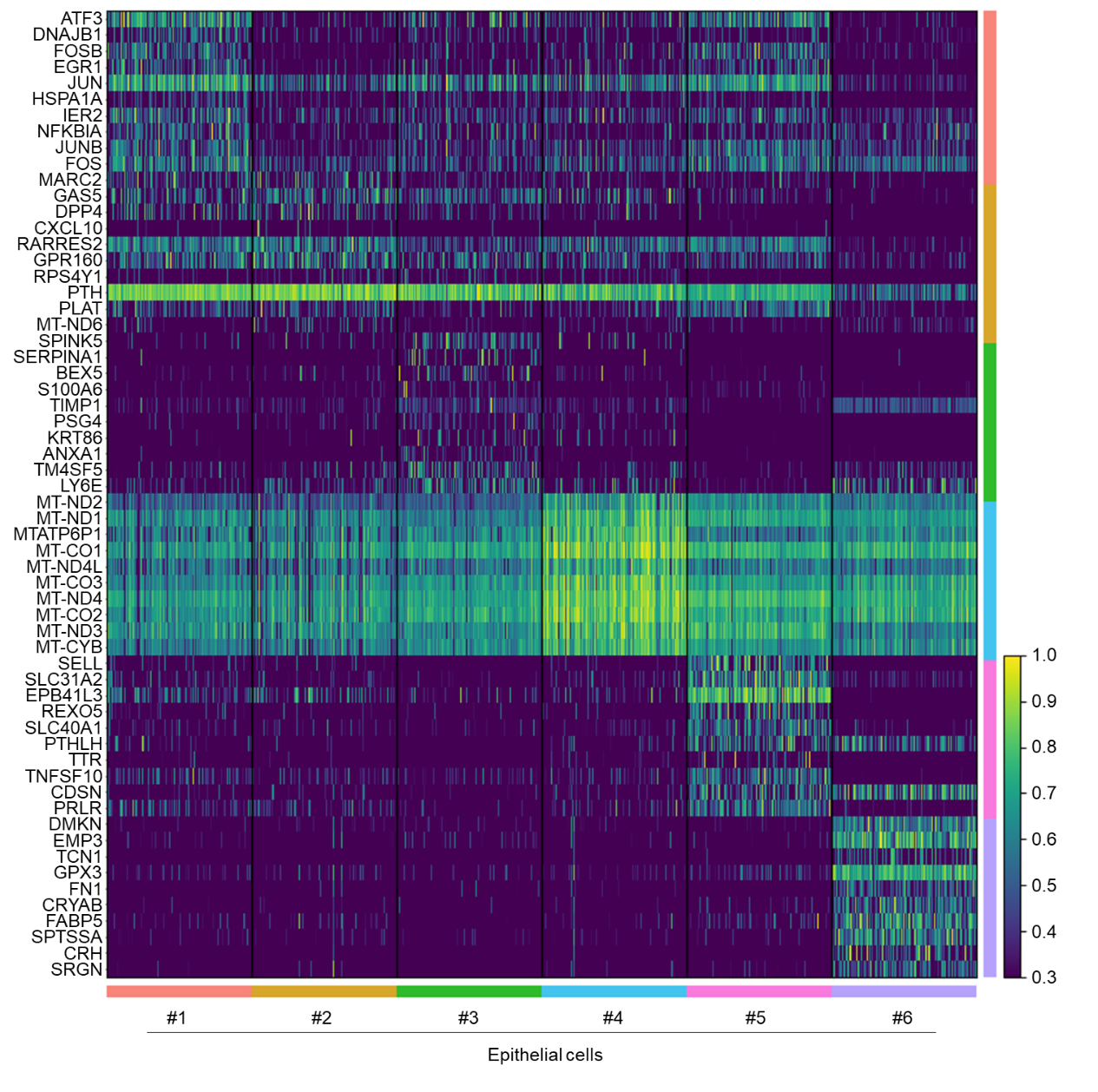
**Supplemental Figure 6. Transcriptomic heterogeneity of parathyroid epithelial cells.** Heatmap displaying the scaled expression of the top 10 DEGs for each of the six identified epithelial subclusters.

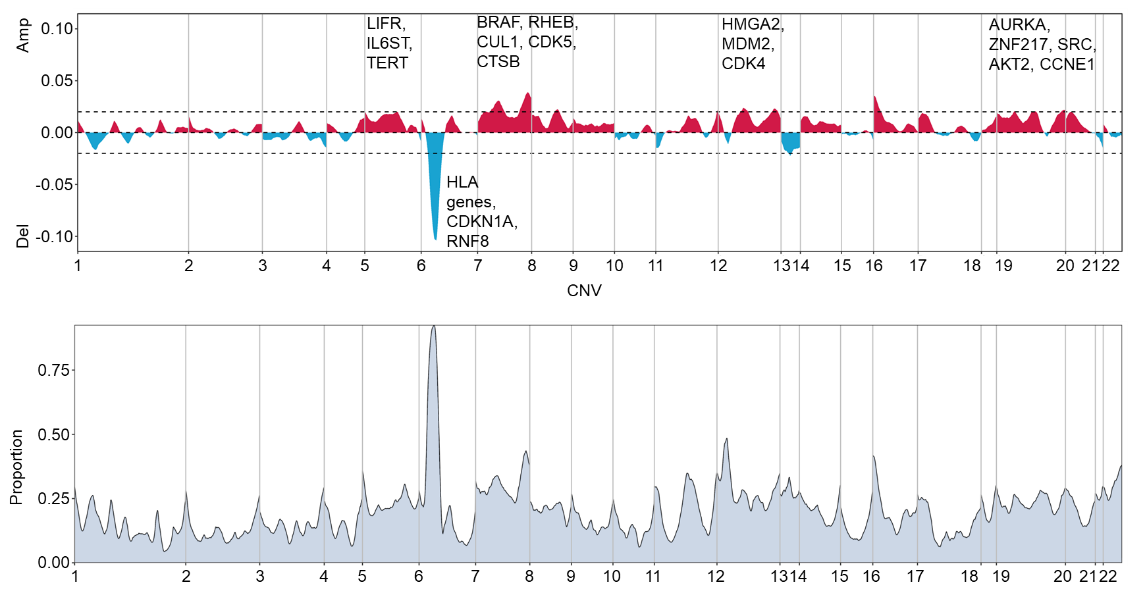

**Supplemental Figure 7. Inferred genomic landscape and key somatic copy number alterations.** Genome-wide copy number variation (CNV) profile inferred from scRNA-seq data across parathyroid epithelial cells. The upper track displays the aggregate CNV score (amplitude), with **red** areas indicating chromosomal **amplifications (Amp)** and **blue** areas indicating **deletions** (Del). Key genes associated with parathyroid tumorigenesis and disease progression located within these significant CNV regions are annotated. The lower track illustrates the proportion of cells harboring the corresponding CNV events across the genome.

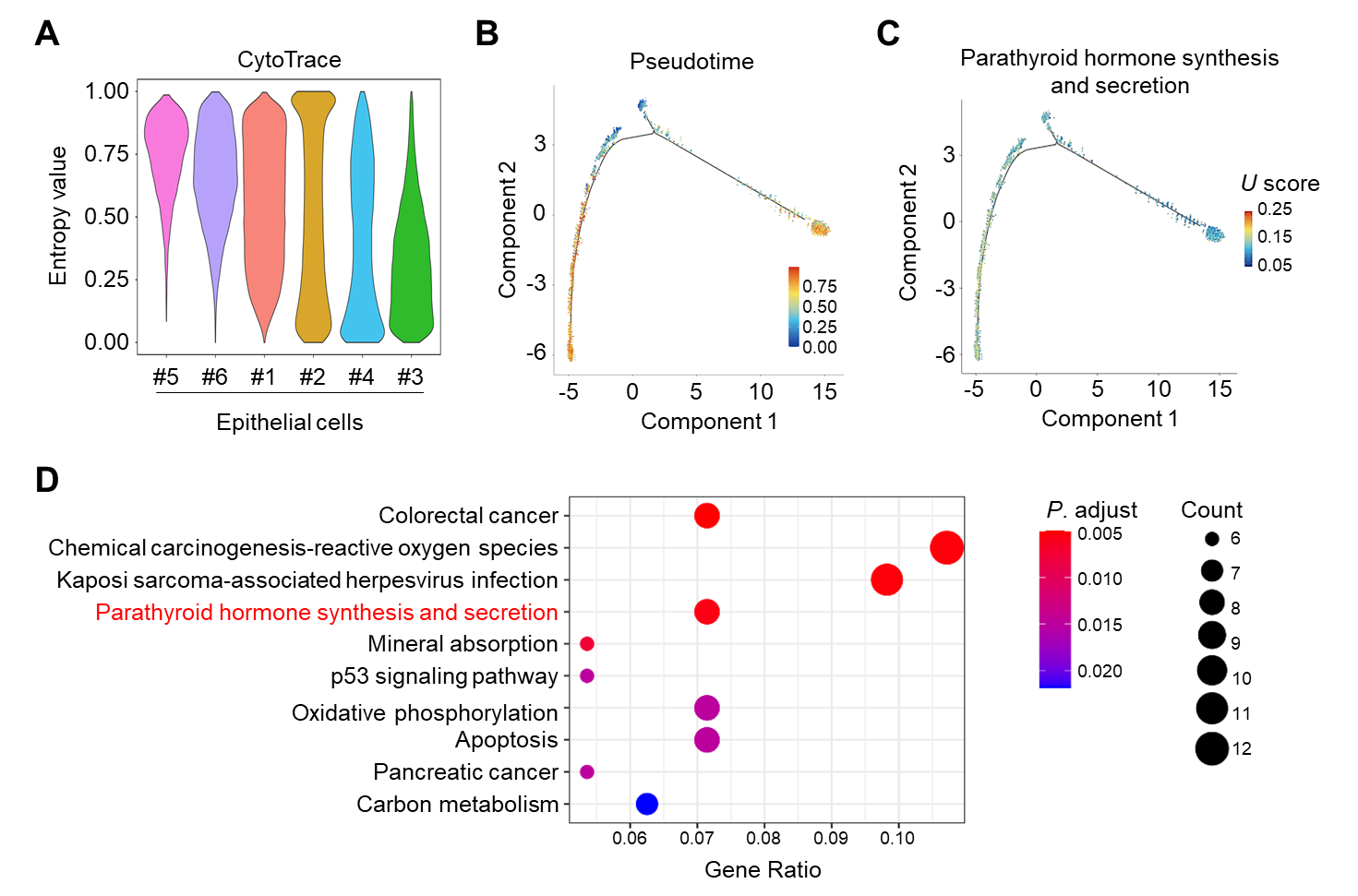

**Supplemental Figure 8. Functional characterization and trajectory analysis of parathyroid epithelial cells. (A)** Violin plots showing the CytoTRACE score (entropy value) across different epithelial subclusters. **(B, C)** Monocle 2 pseudotime trajectory of epithelial cells. Individual cells are colored by their CytoTRACE score (**B**) and parathyroid hormone (PTH) synthesis and secretion signature score (expressed as U score) (**C**). **(D)** Dot plot illustrating the KEGG pathway enrichment analysis of downregulated DEGs for Ep #3 compared to the other epithelial subclusters.

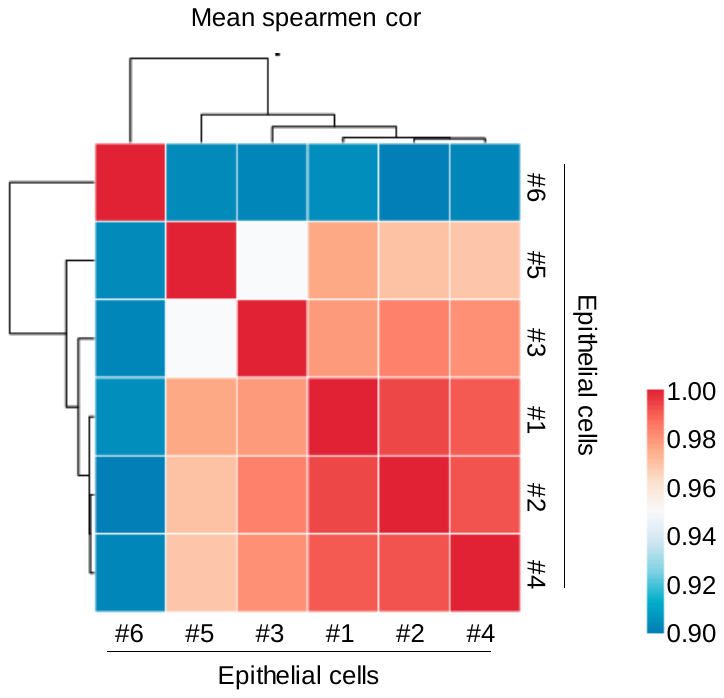

**Supplemental Figure 9. Transcriptomic correlation analysis of epithelial subclusters.** Heatmap illustrating the **Pearson correlation coefficients** between the average gene expression profiles of the six identified epithelial subclusters (Ep #1 – #6). Hierarchical clustering (dendrograms) groups subclusters with similar transcriptomic signatures, revealing potential lineage relationships or functional similarities.

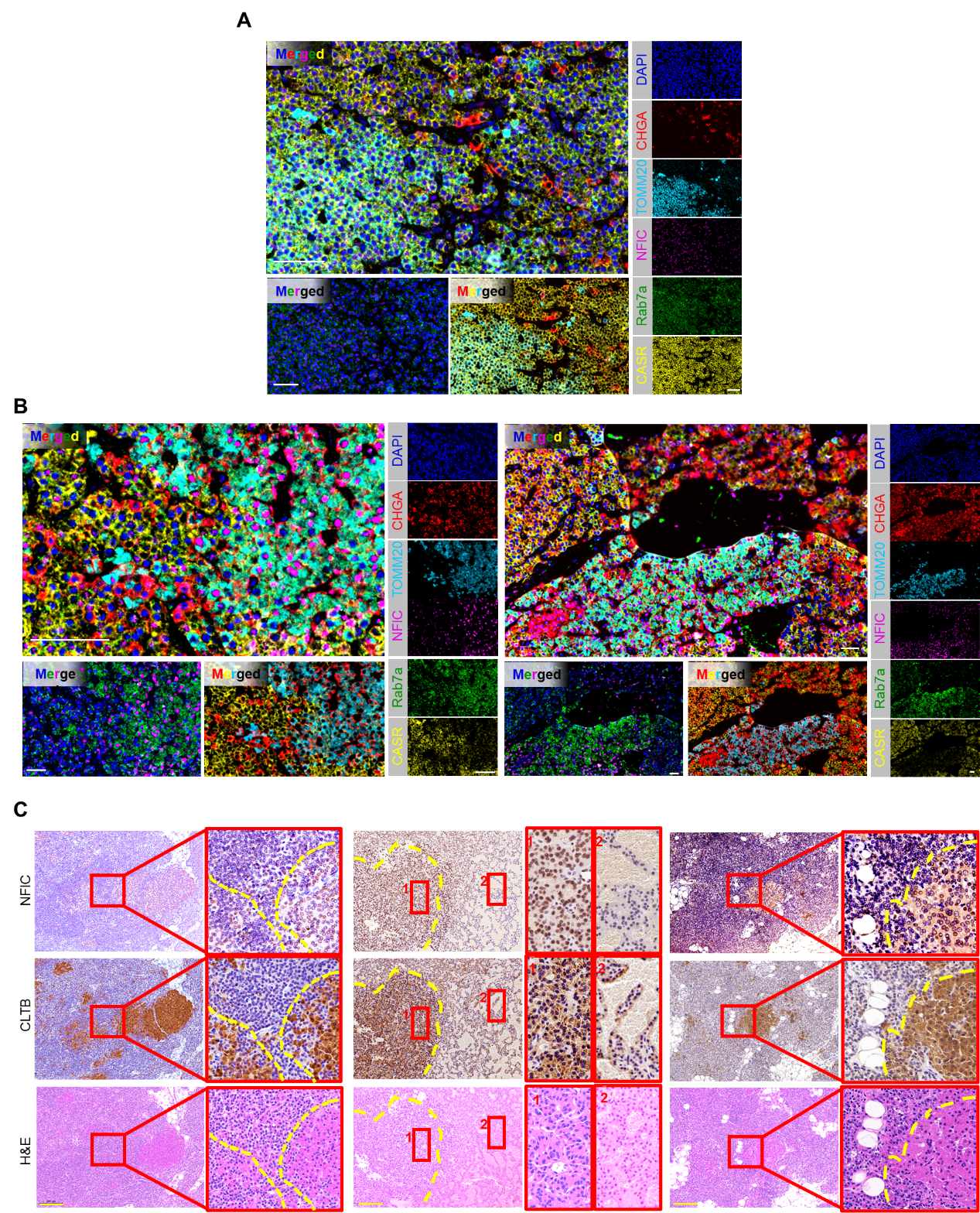

**Supplemental Figure 10. Multiplex immunofluorescence (mIF) characterization of parathyroid cell populations in refractory SHPT.** Representative mIF images of FFPE tissue sections from refractory SHPT patients, stained using Tyramide Signal Amplification (TSA) technology. The panel displays the expression and spatial distribution of CHGA (red), TOMM20 (cyan, mitochondrial marker), NFIC (magenta), Rab7a (green), and CASR (yellow). Nuclei are stained with DAPI (blue). **(A)** Visualization of **chief cells** and **conventional oxyphil cells**. Chief cells are characterized by TOMM20^low^CHGA^high^, while conventional oxyphil cells show TOMM20^high^CHGA^low^. **(B)** Extended for Fig. 5g, providing expanded views of **chief cells (**TOMM20^low^CHGA^high^**), along with** pathological **oxyphil cells (TOMM20^high^CHGA^high^). The latter was** coupled with **elevated NFIC and Rab7a** levels, but notably **reduced membrane CaSR** intensity compared to chief cells. Scale bars: 50 μm. **(C)** Serial section analysis of SHPT tissues using H&E staining and IHC for NFIC and CLTB. Three representative regions are shown. The yellow dashed lines delineate the boundaries between pathological oxyphil nodules (eosinophilic in H&E) and surrounding chief cells. High-magnification insets (red boxes) confirm that NFIC^high^ nuclear staining strictly colocalizes with CLTB^high^ cytoplasmic staining within the deep eosinophilic (oxyphil) regions, whereas adjacent chief cell regions show low expression of both proteins. This spatial correlation supports the hypothesis that NFIC transcriptionally orchestrates the upregulation of the endocytic adaptor CLTB in pathological oxyphil cells. Scale bars: 200 μm.

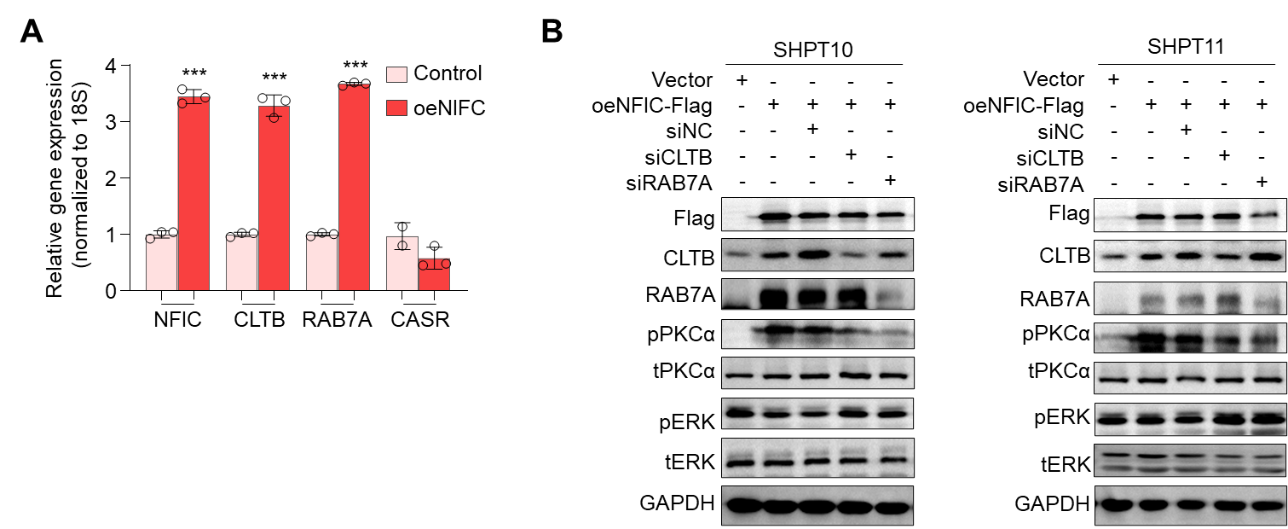

**Supplemental Figure 11. Validation of NFIC-mediated regulation of CaSR and endocytic machinery in primary parathyroid cells.** **(A)** RT-qPCR analysis of relative mRNA expression levels of NFIC, CLTB, RAB7A, and CASR in primary parathyroid cells transfected with an empty vector (control) or an NFIC overexpression plasmid (oeNFIC). Data are presented as mean ± SD (n = 3). Statistical significance was determined using a two-tailed Student’s t-test. ***, *P* < 0.001. **(B)** Western blotting analysis of key proteins involved in the endocytic and lysosomal degradation pathway. Primary parathyroid cells were subjected to either NFIC overexpression or siRNA-mediated CLTB/Rab7a knockdown. GAPDH served as the loading control.
