## Supplemental Tables for "NFIC-mediated CaSR endocytosis defines a hyperactive TOMM20^high^CHGA^high^ oxyphil cell state as a pathological driver of autonomous secondary hyperparathyroidism"

Qi Yang *et al*.

**Supplemental Table 1. Clinical characteristics of patients enrolled in this study.**

| **Patient ID** | **Experimental Usage** | **Age (yr)** | **Sex** | **Primary Disease** | **CKD Stage** | **Pre-surgical intervention** | | | | | **Pre-op iPTH (pg/mL) [15.0-65.0]** | **Pre-op Ca (mmol/L) [2.11-2.52]** | **Serum P (mmol/L) [0.85-1.51]** | **ALP (U/L) [45-125]** |
| --- | --- | --- | --- | --- | --- | --- | --- | --- | --- | --- | --- | --- | --- | --- |
|  |  |  |  |  |  | **Dialysis Modality** | **Kidney Transplant** | **Calci-mimetics** | **Active Vit D** | **Phosphate Binders** |  |  |  |  |
| PC1 | scRNA-seq | 73 | M | PC | NA | NA | NA |  |  |  | 2154.0 | 4.73 | 1.18 | 879 |
| PA1 | scRNA-seq | 36 | M | PA | NA | NA | NA |  |  |  | 95.4 | 2.37 | 1.14 | 114 |
| PA2 | scRNA-seq | 51 | F | PA | NA | NA | NA |  |  |  | 1122.0 | 3.49 | 0.70 | 322 |
| SHPT1 | scRNA-seq | 33 | M | CGN | 3 | NA | Yes |  |  | √ | 1665.0 | 2.19 | 1.85 | 352 |
| SHPT2 | scRNA-seq | 61 | M | HN | 4 | PD | No | √ | √ | √ | 1665.0 | 1.83 | 1.50 | 129 |
| SHPT3 | scRNA-seq | 49 | F | HN | 4 | PD | No |  | √ |  | 1184.3 | 2.15 | 1.49 | 420 |
| SHPT4 | scRNA-seq | 62 | F | PKD | 5 | HD | No | √ |  |  | 2517.0 | 2.42 | 0.60 | 107 |
| SHPT5 | Primary culture | 39 | M | CGN | 5 | HD | No |  |  | √ | 2008.8 | 2.44 | 1.94 | 218 |
| SHPT6 | Primary culture | 36 | M | CGN | 5 | HD | No |  | √ |  | 1228.9 | 1.20 | 0.60 | 364 |
| SHPT7 | Primary culture | 42 | F | CGN | 5 | HD | No | √ |  |  | 3233.0 | 2.54 | 1.99 | 660 |
| SHPT8 | Primary culture | 38 | F | CGN | 5 | HD | No |  | √ | √ | 1508.0 | 2.10 | 2.40 | 474 |
| SHPT9 | Primary culture | 55 | F | DN | 5 | HD | No | √ |  | √ | 3024.0 | 2.61 | 1.36 | 811 |
| SHPT10 | Primary culture | 48 | M | HN | 5 | HD | No |  |  | √ | 1203.0 | 2.03 | 1.57 | 108 |
| SHPT11 | Primary culture | 36 | F | CGN | 5 | HD | No |  |  | √ | 1184.0 | 3.02 | 3.00 | 361 |
| SHPT12 | Primary culture | 73 | F | HN | 5 | HD | No |  |  | √ | 1150.0 | 2.97 | 0.80 | 248 |
| SHPT13 | Primary culture | 31 | F | CGN | 5 | HD | No |  |  |  | 1922.0 | 2.47 | 0.53 | 244 |
| SHPT14 | Primary culture | 41 | M | HN | 5 | HD | No |  |  |  | 2050.4 | 2.69 | 0.78 | 395 |
| SHPT15 | Primary culture | 37 | F | UNK | 3 | NA | YES |  |  |  | 1349.6 | 2.97 | 0.43 | 173 |
| SHPT16 | Primary culture | 40 | M | UNK | 5 | HD | No | √ |  |  | 2100.1 | 2.63 | 1.23 | 163 |
| SHPT17 | Primary culture | 36 | M | CGN | 5 | HD | No |  | √ |  | 2200.3 | 2.07 | 1.93 | 273 |
| SHPT18 | Primary culture | 62 | F | UNK | 5 | HD | No |  | √ |  | 1700.3 | 1.80 | 0.71 | 162 |
| SHPT20 | Primary culture | 37 | M | CGN | 5 | PD | No |  |  |  | 1264.4 | 2.52 | 1.77 | 159 |
| SHPT22 | Primary culture | 48 | M | UNK | 5 | HD | No | √ | √ |  | 1781.8 | 1.75 | 0.97 | 236 |
| SHPT23 | Primary culture | 63 | F | DN | 5 | HD | No |  |  |  | 1800.6 | 2.68 | 0.85 | 405 |
| SHPT24 | Primary culture | 46 | F | CGN | 5 | HD | No | √ | √ |  | 1433.0 | 1.61 | 2.45 | 143 |
| SHPT25 | Primary culture | 39 | F | CGN | 5 | HD | No | √ | √ |  | 2287.4 | 2.12 | 2.19 | 298 |
| SHPT26 | Primary culture | 35 | M | HN | 5 | HD | No | √ | √ | √ | 805.9 | 1.98 | 1.68 | 222 |
| SHPT27 | Primary culture | 57 | F | HN | 3 | NA | YES |  | √ | √ | 1000.6 | 2.21 | 2.41 | 131 |
| SHPT28 | PDX model | 69 | M | DN | 5 | HD | No | √ |  |  | 929.3 | 2.54 | 0.73 | 70 |
| SHPT29 | PDX model | 39 | F | CGN | 5 | HD | No |  |  |  | 1153.7 | 2.53 | 1.06 | 108 |
| SHPT30 | PDX model | 30 | F | CGN | 5 | HD | No |  |  |  | 1003.4 | 3.03 | 0.80 | 340 |
| SHPT31 | PDX model | 56 | M | HN | 5 | PD | No |  |  |  | 881.4 | 2.77 | 0.48 | 289 |
| SHPT32 | PDX model | 33 | M | CGN | 5 | HD | No |  |  |  | 684.8 | 2.74 | 0.55 | 246 |

Abbreviation: M/F: Male/Female; PC: parathyroid carcinoma; PA: parathyroid adenoma; SHPT: secondary hyperparathyroidism; Primary Disease: CGN, chronic glomerulonephritis; HN, hypertensive nephrosclerosis; PKD, polycystic kidney disease; DN, diabetic nephropathy; UNK, unknown; CKD: chronic kidney disease; Dialysis Modality: HD, hemodialysis; PD, peritoneal dialysis; iPTH: intact parathyroid hormone; Ca: serum calcium; P: serum phosphorus; ALP: alkaline phosphatase; NA: not applicable.

**Supplemental Table 2. List of cell type annotations.**

| **Cell type** | **Markers** |
| --- | --- |
| Epithelial cells | EPCAM, KRT19, KRT18 |
| Endothelial cells | PECAM1, VWF, CDH5 |
| Mural cells | ACTA2, RGS5, PDGFRB |
| Immune cells | PTPRC |
| Fibroblasts | DCN, COL1A2, COL1A1 |
| T and NK | CD3D, CD3G, KLRF1 |
| Mast Cells | TPSAB1, TPSB2, CPA3 |
| Macrophages | C1QA, MRC1, CD163 |
| Monocytes | CD14, VCAN, FCN1 |
| cDC2 | CD1C, CD1E, FCER1A |
| AECs | GJA5, DKK2, BMX |
| Tip | CXCR4, KCNE3, ESM1, ANGPT2 |
| VECs | ACKR1, SELP, NR2F2, SELE |
| Cap ECs | RGCC, CA4, EDNRB, FCN3 |

| **Functional State / Subpopulation** | **Gene Signature (Marker Genes)** | **Analysis Method** |
| --- | --- | --- |
| Tissues repair /angiogenesis | *EREG*, *AREG*, *VEGFA*, *THBS1*, *CD83* | GSVA scoring |
| Pro-inflammatory | *IL1B*, *CXCL8*, *CXCL1*, *CXCL2*, *CXCL3*, *CCL3*, *CCL4*, *NAMPT*, *NFKBIA* | GSVA scoring |
| Nerve-associated macrophage | *P2RY12*, *P2RY13*, *CX3CR1*, *TMEM119*, *C3*, *BIN1*, *HEXB* | GSVA scoring |
| Perivascular resident | *LYVE1*, *FOLR2*, *MRC1*, *SELENOP*, *CD163*, *F13A1*, *SLC40A1*, *STAB1*, *IGF1* | GSVA scoring |
| Non-classical monocyte | *FCGR3A*, *LST1*, *CDKN1C*, *HAMP*, *LILRB2*, *SMIM25*, *MS4A7*, *VMO1* | GSVA scoring |
| Osteoclast-like | *ACP5*, *LIPA*, *CTSD*, *FABP5*, *LPL*, *GPNMB*, *LGALS3*, *TREM2*, *APOE* | GSVA scoring |
| Proliferation | *MKI67*, *TOP2A*, *CDK1*, *PCNA*, *CCNB1*, *CDC20*, *TYMS* | GSVA scoring |

**Supplemental Table 3. Curated gene signatures used for macrophage subpopulations.**

**Supplemental Table 4**. **The sequences of siRNAs used in this study.**

| **siRNAs** | **Sense (5’-3’)** | **Antisense (5’-3’)** |
| --- | --- | --- |
| siNC | UUCUCCGAACGUGUCACGUTT | AAACGTGACACGTTCGGAGAATT |
| siCLTB-1 | GAUGCUGCAUCUAAGGUCATT | UGACCUUAGAUGCAGCAUCTT |
| siCLTB-2 | CGGUGCUCAUGUCCCUGAATT | UUCAGGGACAUGAGCACCGTT |
| siCLTB-3 | GUAGAGAAGAACAAGAUCATT | UGAUCUUGUUCUUCUCUACTT |
| siRAB7A-1 | GGAUGACCUCUAGGAAGAATT | UUCUUCCUAGAGGUCAUCCTT |
| siRAB7A-2 | GUACAACGAAUUUCCUGAATT | UUCAGGAAAUUCGUUGUACTT |
| siRAB7A-3 | GCCACAAUAGGAGCUGACUUUTT | AAAGUCAGCUCCUAUUGUGGCTT |

**Supplemental Table 5. List of primary antibodies used in this study.**

| **Antigen** | **Brand** | **Cat No.** | **Application** |
| --- | --- | --- | --- |
| CaSR | Proteintech | 19125-1 | IHC, ICC, mIF (FFPE) |
| CLTB | Zen Bio | 126146 | IHC, ICC, mIF (FFPE), Western Blot |
| Rab7A | CST | 95764 | ICC, mIF (FFPE), Western Blot |
| CHGA | Zen Bio | 381686 | mIF (FFPE) |
| TOMM20 | Zen Bio | 382451 | mIF (FFPE) |
| NFIC | Proteintech | 16399-1 | IHC, ICC, mIF (FFPE), Western Blot |
| Ki67 | Abcam | ab15580 | IHC |
| AP2B1 | Proteintech | 15690-1-AP | ICC |
| LAMP1 | Invitrogen | 14-1071-82 | ICC |

| **Targets** | | **Oligonucleotides (5’- 3’)** | **Application** |
| --- | --- | --- | --- |
| PTH | Forward | CTCAGCATCAGCTACTAACATACC | RT-qPCR |
|  | Reverse | CACTCACAGATCTCTTCTTAACAG | RT-qPCR |
| CASR | Forward | GCAGAATGAAAGGCATCACAG | RT-qPCR |
|  | Reverse | AGAAGACGTGGTGTTTGAATCTC | RT-qPCR |
| CLTB | Forward | GATCAACAACCGGATCGCTG | RT-qPCR |
|  | Reverse | GGGTTGAAGTCACATAGCTGG | RT-qPCR |
| RAB7A | Forward | GCTTCTGTCCTCCGTTTAGTC | RT-qPCR |
|  | Reverse | TTCCCGACTCCAGAATCTCC | RT-qPCR |
| NFIC | Forward | GACCTCGCCTTCCTACTCTC | RT-qPCR |
|  | Reverse | GACCTTTGCTATCCCAGATACC | RT-qPCR |
| RPS18 | Forward | CAGCCACCCGAGATTGAGCA | RT-qPCR |
|  | Reverse | TAGTAGCGACGGGCGGTGTG | RT-qPCR |
| CLTB promoter | #1 Forward | ACTATATTGACCAGGCTGGTCG | ChIP-qPCR |
|  | #1 Reverse | AAGCAGCAGGATACAAAGCCA | ChIP-qPCR |
|  | #2 Forward | GACCAGCCCAGTTAATATAGCGA | ChIP-qPCR |
|  | #2 Reverse | GTTTCTTTGCATCGTCTCCC | ChIP-qPCR |
|  | #3 Forward | CTGATGACTTTGGCTTCTTCTC | ChIP-qPCR |
|  | #3 Reverse | GACGGTTAGGGCATTCTAAATCAC | ChIP-qPCR |
|  | #4 Forward | CTGGGCTCAAGTGATCCTCTC | ChIP-qPCR |
|  | #4 Reverse | GCTGTCTGCTATTTCTAATGTGG | ChIP-qPCR |
|  | #5 Forward | CTCTGAATGCCTCCTCCCTG | ChIP-qPCR |
|  | #5 Reverse | CCTCGTCGTTCTCTATGCCT | ChIP-qPCR |
| RAB7A promoter | #1 Forward | CAGGAGTTCAAGACTGCCTG | ChIP-qPCR |
|  | #1 Reverse | AGTGCAGTGGCATAATCAGGG | ChIP-qPCR |
|  | #2 Forward | GAAGAGCTTTCCAGGTAGAG | ChIP-qPCR |
|  | #2 Reverse | CCACTCTTTCTCTGCTTTCAC | ChIP-qPCR |
|  | #3 Forward | GACTGCCTTTCTCTGGTCTG | ChIP-qPCR |
|  | #3 Reverse | GCAATGTGAGGCAACCATGAC | ChIP-qPCR |
|  | #4 Forward | AAGTGCCTATTTCTCCTGTC | ChIP-qPCR |
|  | #4 Reverse | TCTGTCTGATGCCAATATGTG | ChIP-qPCR |
|  | #5 Forward | TGTTTGTAGCACGAGATCCAGG | ChIP-qPCR |
|  | #5 Reverse | GGCTTTATGGCCAAGACTCC | ChIP-qPCR |

**Supplemental Table 6. List of primers used in this study.**
